# Structural basis for ATP-driven double-ring assembly of the human mitochondrial Hsp60 chaperonin

**DOI:** 10.1101/2025.10.04.680452

**Authors:** Igor Tascón, Jorge P. López-Alonso, Yoel Shkolnisky, David Gil-Cartón, Jesús Vilchez-Garcia, Alberto G. Berruezo, Yacob Gómez-Llorente, Radhika Malik, Fady Jebara, Malay Patra, Joel A. Hirsch, Abdussalam Azem, Iban Ubarretxena-Belandia

## Abstract

The ATP-driven mHsp60:mHsp10 chaperonin system assists protein folding within the mitochondrial matrix of human cells. Substrate protein folding has been proposed to occur through interconnected single- and double-ring pathways. In the absence of nucleotide, mHsp60 exists in equilibrium between free protomers and heptameric single rings, while the formation of double rings requires ATP. Here, we present cryo-electron microscopy structures of mHsp60 in the apo state, bound to ATP, and bound to ATP in complex with the cochaperonin mHsp10. ATP binding to single-ring apo mHsp60_7_ triggers coordinated conformational changes in the intermediate and apical domains, resulting in a highly dynamic apical region within the ring. Extensive inter-subunit rearrangements flatten the equatorial surface of each ring, thereby enabling inter-ring contacts that stitch the rings together to form double-ring mHsp60_14_. Collectively, these structures define the structural basis of ATP-driven double-ring assembly of a human mitochondrial chaperonin responsible for maintaining mitochondrial protein homeostasis.

## INTRODUCTION

Driven by ATP, the human mitochondrial group I chaperonin mHsp60 (also known as *HSPD1*) and its co-chaperonin mHsp10 (*HSPE1*) assist the folding of nearly half of all nuclear-encoded proteins imported into the mitochondrial matrix and correct misfolded polypeptides generated under proteostatic stress^1–5^. Genetic studies underscore the essential role of mHsp60. Deletion of *HSPD1* is lethal in yeast^2^, and its inactivation in mice results in embryonic lethality^6^. Defects in mHsp60-mediated protein folding are associated with inherited neurodegenerative diseases^7,8^, while dysregulation of its function has been linked to inflammatory signaling^9,10^, apoptosis^11,12^, and tumor biology, where it is both a prognostic biomarker^13,14^ and a therapeutic target^15,16^.

Each mHsp60 protomer consists of an equatorial domain responsible for ATP binding and hydrolysis, an intermediate domain containing hinge regions that mediate conformational changes, and an apical domain that engages substrate proteins (SPs) and the cochaperonin mHsp10^17,18^. Seven mHsp60 protomers assemble into heptameric rings, which are capped by dome-shaped heptamerice mHsp10 lids to form nano-cages for SPs to fold in confinement^17,18^. While much of our mechanistic understanding of chaperonins comes from the bacterial GroEL:GroES system, biochemical and structural studies have revealed important differences between GroEL and mHsp60^17,19–21^, despite their high-sequence identity (Fig. S1).

In the absence of nucleotide apo GroEL readily forms stable back-to-back heptameric double rings^22,23^. ATP-dependent allosteric transitions drive the formation of asymmetric bullet-shaped complexes, where only one of the two rings in double-ring GroEL is capped by its cochaperonin GroES^24^, and symmetric American football-shaped complexes, where both rings are capped^25–27^. In *E. coli* these double-ring GroEL:GroES asymmetric and symmetric complexes are functionally linked^28^.

In contrast, mHsp60 has been proposed to fold SP through interconnected single- and double-ring pathways^17,19–21^. Apo mHsp60 exists in equilibrium between free protomers and heptameric single rings, with the formation of double rings requiring ATP^17,18,29^. In the presence of ATP and mHsp10, both double-ring football complexes and single-ring half-football complexes have been observed^18,29–31^. We reported the first crystal structure of mHsp60 using the E321K chaperonin variant, which stabilized its association with mHsp10^18,32^, and later determined the crystal structure of wild-type (WT) mHsp60 bound to mHsp10 and the ATP ground-state mimic ADP:BeF_3_^17^. Both crystal structures captured double-ring football assemblies, whereas single-ring half-football complexes could not be crystallized. Cryo-electron microscopy (cryo-EM) enabled structure determination of ADP-bound half-football and football assemblies imaged together from the same specimen, providing early structural evidence for the co-existence of single- and double-ring mHsp60:mHsp10 complexes^17^. The cryo-EM structures confirmed that half-football complexes mirror the architecture of one half of the football assembly and showed that double-ring stability is mediated by a two-fold symmetric hydrogen bond between S464 residues of opposing rings reinforced by a salt bridge^17^. Remarkably, engineered interface variants producing obligated single- or double-ring assemblies retained folding activity in vitro, and bacterial complementation assays demonstrated that the single-ring variant was as efficient as WT mHsp60^17^. More recent cryo-EM and molecular dynamics studies have captured multiple conformational states of both single- and double-ring mHsp60:mHsp10 complexes, providing further support for a pathway involving football and half-football complexes^33–37^.

However, the structural basis for why mHsp60 fails to assemble into double rings in the absence of nucleotide remains unclear. Here, we present cryo-EM structures of purified WT mHsp60 in its apo state, bound to ATP, and bound to both ATP and mHsp10, which together uncover the structural basis of ATP-driven double-ring assembly of the human mitochondrial chaperonin.

## RESULTS

### Cryo-EM structures of ATP-bound mHsp60 complexes

In the present work, we investigated the structural effects of ATP binding, for which we relied on comparing cryo-EM structures determined from the same protein preparation. Cryo-EM imaging (Fig. S2) and 3D reconstruction (Fig. S3) were carried out using recombinant WT human mHsp60 and mHsp10^17,38^. Reconstructions were calculated with D7 symmetry for double-ring complexes and C7 symmetry for single rings, with C1 reconstructions without symmetry imposition generated for reference (Fig. S3). Table 1 summarizes structure determination statistics.

**Table 1.**
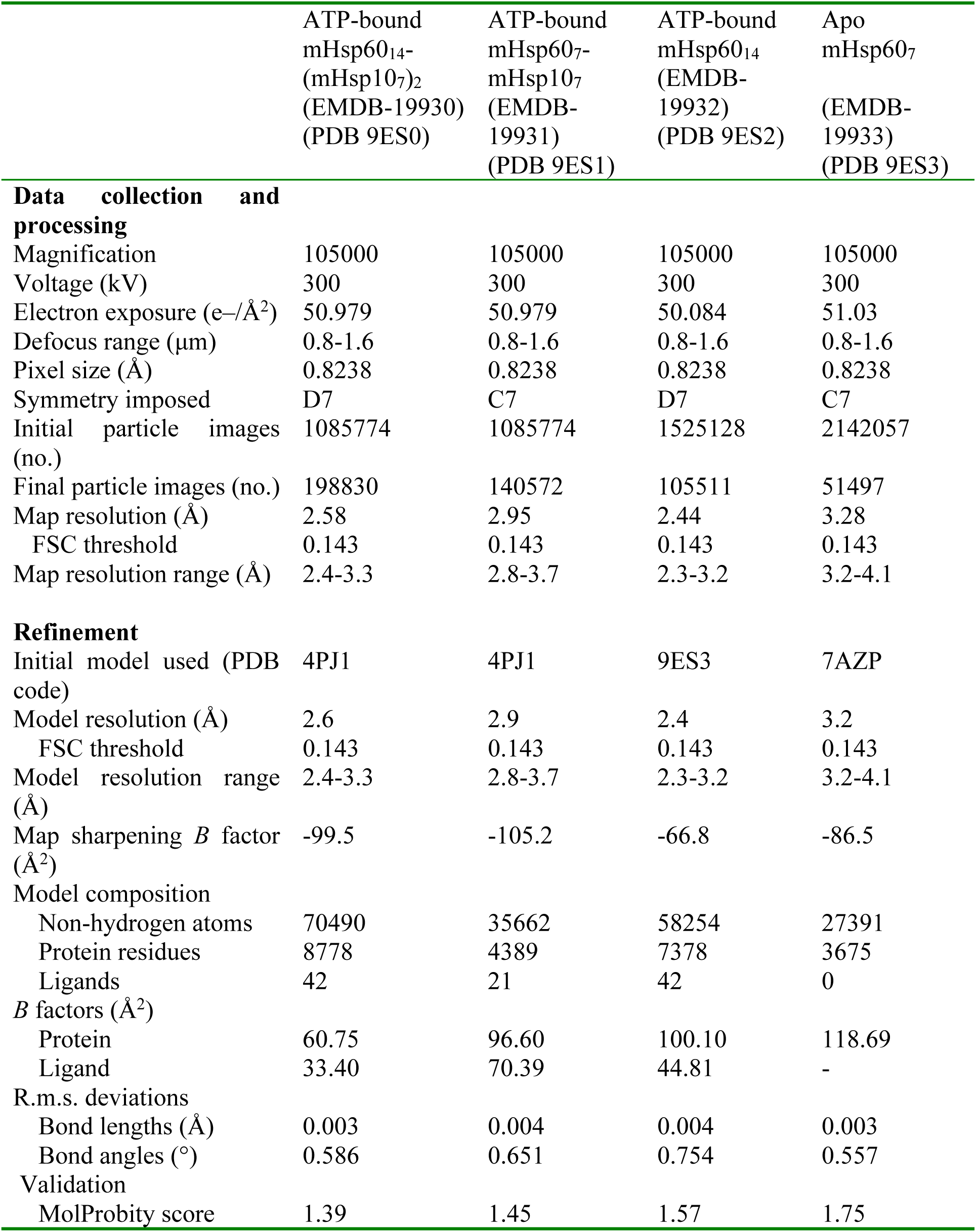
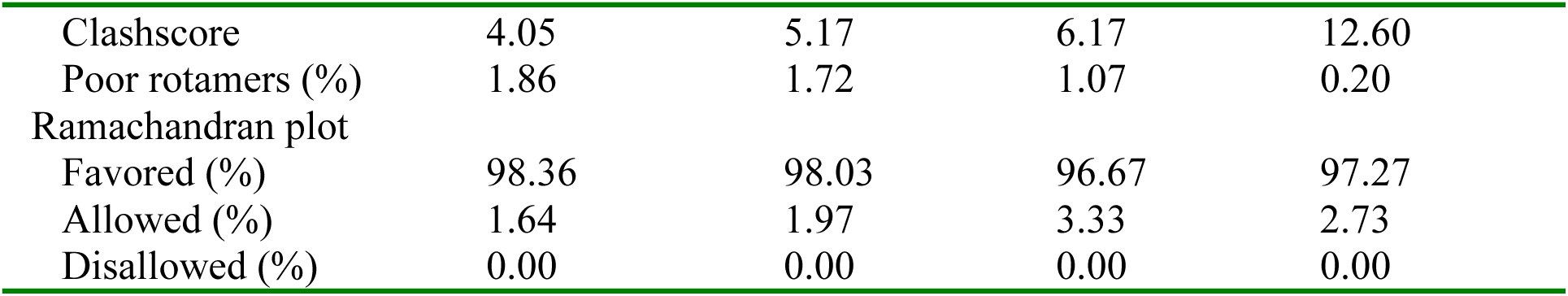
Cryo-EM data collection, refinement and validation statistics.

At a nominal resolution of 3.28 Å with C7 symmetry imposed (Fig. S3A) the cryo-EM map of apo mHsp60 shows protomers arranged in a single heptamer ring (Fig. S4, map of apo mHsp60_7_ shown in red). In line with biochemical experiments^17,29^, along with single-rings we also observed mHsp60 protomers in the cryo-EM images, but not double rings (Fig. S2, left panel). Each subunit in the heptameric single-ring structure (Fig. 1, PDB ID 9ES3) comprises an equatorial domain (residues 1–137; 411–526) responsible for nucleotide binding and inter-subunit interactions, an apical domain (residues 192–374) involved in SP and co-chaperonin binding, and an intermediate domain (residues 138– 191; 375–411) with two hinge regions (Fig. S5). The nucleotide-binding pockets are empty (Fig. S4), confirming this single-ring structure as an apo species, resembling recently reported cryo-EM structures of nucleotide-free mHsp60^33,34,36^.

**Figure 1.**
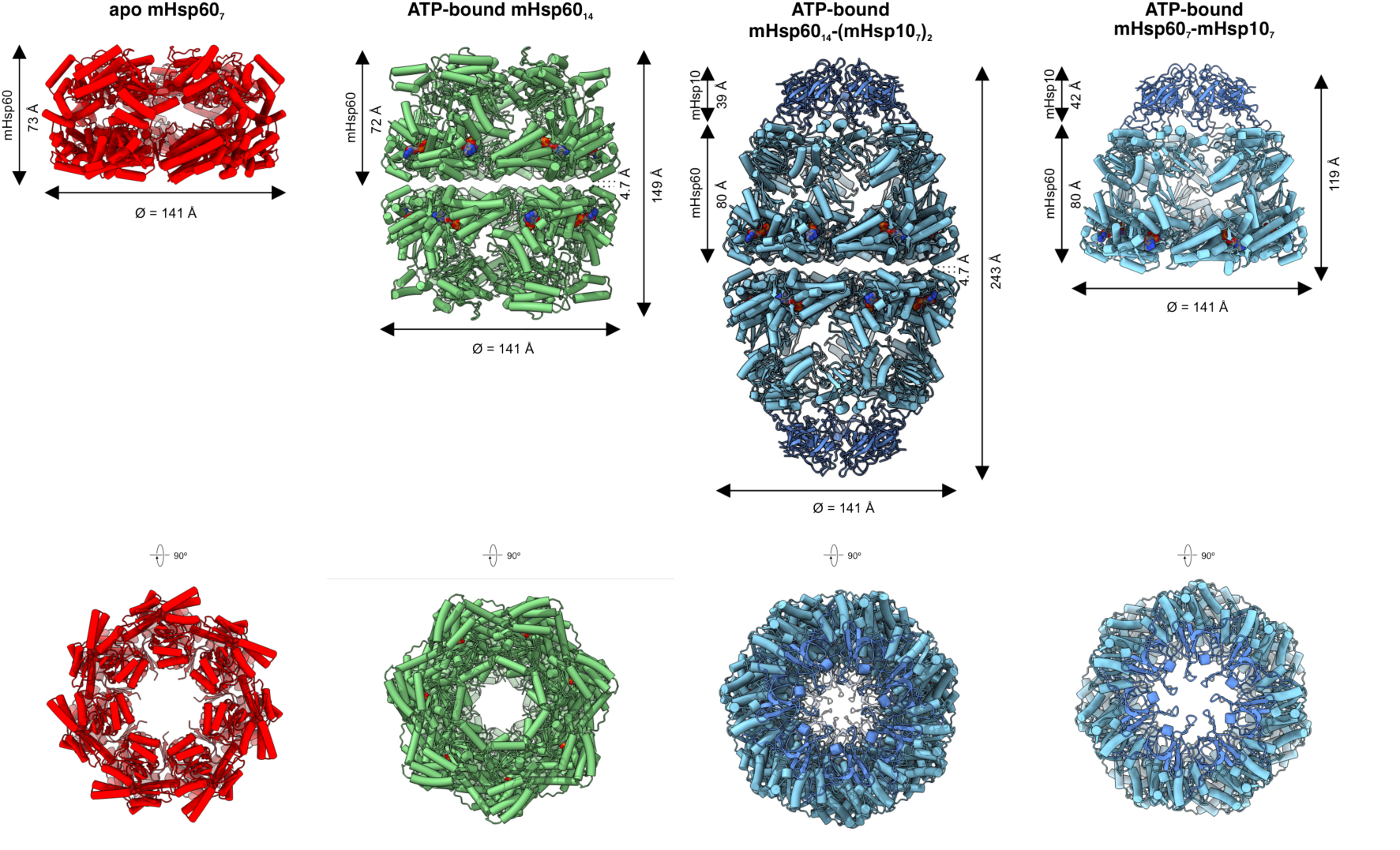
Cryo-EM structures of mHsp60 complexes. Cartoon depiction of the structures of apo mHsp60_7_ (red), ATP-bound mHsp60_14_ (green), ATP-bound mHsp60_14_-(mHsp10_7_)_2_ and ATP-bound mHsp60_7_-mHsp10_7_ (mHsp60 subunits in cyan and mHsp10 subunits in blue). The structures are shown in side (top) and top view (bottom). Space-filling representations depict the nucleotides. Dimensions of the complexes are respectively indicated, using symmetrical Cα to define limiting planes.

To prevent nucleotide hydrolysis and capture ATP-bound states, we adapted the specimen preparation strategy employed for the ADP-bound complexes^17^, with the key modification that vitrification was carried out immediately after mixing mHsp60 with fresh ATP, omitting the 30 min. pre-incubation step used in the ADP dataset. In the presence of ATP both single- and double-ring assemblies were observed (Fig. S2, center panel). The scarcity and preferred top view orientation of the single-rings precluded their structure determination, leaving their nucleotide state unresolved. For the double-ring complexes, a 2.44 Å nominal resolution map obtained with D7 symmetry (Fig. S3B) captured mHsp60 heptameric rings stacked back-to-back (Fig. S4, ATP-bound mHsp60_14_ shown in green). Intermediate and equatorial domains were well resolved, while apical densities were weak and discontinuous. Gaussian filtering of the ATP-bound cryo-EM map with a 1.47 standard deviation in Chimera^39^ allowed visualization of the apical region density (Fig. S4, ATP-bound mHsp60_14_ filtered map). A comparable poorer resolution of the apical domains in the ATP-bound mHsp60 species has been recently observed for a V72I mHsp60 variant displaying increased heptamerization^33^. In that case a model for ATP-bound double-ring mHsp60^V72I^ was built by docking the mHsp60 double-ring from our ADP-bound mHsp60:mHsp10 football structure^17^, followed by refinement. In this case, we built de novo the structure of the equatorial and intermediate domains based on the high-resolution density features of the ATP-bound mHsp60 cryo-EM map in this region, whereas for the apical region we docked the analogous region of apo mHsp60 (PDB ID 9ES3) followed by refinement. In the resulting mHsp60_14_ structure (Fig. 1, PDB ID 9ES2) all subunits are bound to ATP, Mg^2+^ and K^+^ (Fig. S4).

Addition of mHsp10 yielded ATP-bound football complexes (Fig. S2, right panel; Fig. S3C) at 2.58 Å nominal resolution map (Fig. S4, ATP-bound mHsp60_14_-(mHsp10_7_)_2_ shown in light and dark blue). The structure (Fig. 1, PDB ID 9ES0) shows two stacked heptamer rings capped at both ends by dome-shaped heptameric mHsp10 lids. As for the ADP-bound state (PDB 6MRC)^17^, the architecture is football-shaped with RMSD value of 0.453 Å compared to the ADP-bound football structure^17^. All mHsp60 protomers are bound to ATP, Mg^2+^ and K^+^ (Fig. S4; Fig. S6-7) and adopt an open conformation along the molecular symmetry axis with well-resolved densities for the equatorial, intermediate and apical domains. Relative to their placement in ATP-bound mHsp60, the apical domains rise by 83° and rotate by 90° clockwise (view from within the assembly) in ATP-bound mHsp60-mHsp10 (Fig. S8, right panel; Movie S1). This brings the hydrophobic surface of helices H and I in contact with the mobile loop of each mHsp10 protomer (Fig. S9), which adopts the canonical seven-strand β-barrel structure and exposes a mobile loop that mediates the interaction with helices H and I of the mHsp60 apical domains.

We also observed the characteristic single-ring mHsp60-mHsp10 half-football complex^17^ in the cryo-EM images (Fig. S2, right panel). Independently of the football complexes, we determined their structure at 2.95 Å nominal resolution with C7 symmetry (Fig. S3C; Fig. S4, ATP-bound mHsp60_7_-mHsp10_7_ shown in light and dark blue). This structure (Fig. 1, PDB ID 9ES1) comprises a single-heptameric mHsp60 ring, with all nucleotide binding sites occupied by ATP, Mg^2+^ and K^+^, capped by a single mHsp10 lid. RMSD value of 0.590 Å compared to ADP-bound mHsp60_7_-mHsp10_7_ (PDB 6MRD)^17^, indicates minor overall changes upon ATP hydrolysis.

### ATP-triggered conformational changes in the intermediate and equatorial domains

In the presence of ATP, all subunits in the cryo-EM structures, whether arranged as single- or double-rings, display nucleotide-binding sites occupied by ATP, Mg²⁺, and K⁺ with conserved stereochemistry (Fig. S4). ATP can be confidently distinguished from ADP in these sites (Fig. S6), both in symmetry-imposed and unsymmetrized reconstructions (Fig. S7). Multiple contacts anchor ATP tightly to each nucleotide-binding site (Fig. 2A; Fig. S10). Its adenine moiety stacks against P33 and I494, while the N6 atom forms a hydrogen bond with D480. The side-chain of the catalytic residue D399 activates the catalytic water (number 50) for nucleophilic attack and interacts with the γ-phosphate of ATP. D52 is also near the catalytic water. The metal ion Mg^2+^ is octahedrally coordinated (Fig. 2A), with the ATP triphosphate moiety providing three ligands. Three additional ligands are provided by the side chain of D87 and two water molecules (numbers 148 and 210). K^+^ exhibits distorted polyhedral coordination involving two phosphates, the backbone carbonyls of T30 and K51, the side chain of T90, and a water molecule (number 74). Comparison of the nucleotide-binding sites in mHsp60₁₄ and mHsp60_14_-(mHsp10_7_)_2_ (Fig. 2A; Fig. S11A) indicates that mHsp10 binding does not alter the ATP/Mg²⁺/K⁺ coordination network. By contrast, comparison with the ADP-bound mHsp60:mHsp10 football complexes^17^ reveals a marked loss of nucleotide interactions upon ATP hydrolysis (Fig. S11B).

**Figure 2.**
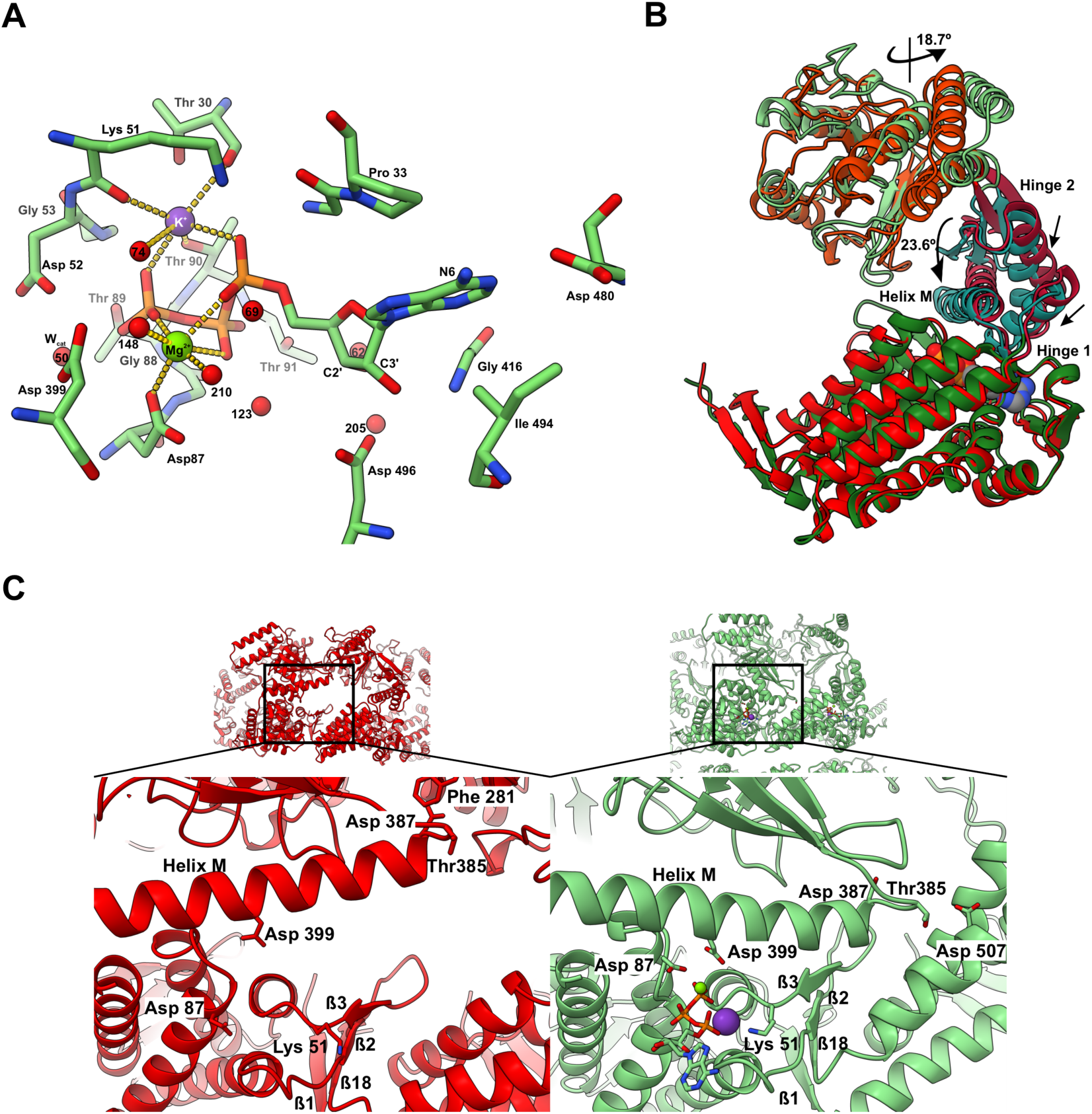
Conformational changes upon ATP binding. (**A**) Stereochemistry of the ATP-bound mHsp60_14_ nucleotide binding site. Close-up focusing on the coordination of the metal ions. The residues interacting with ATP, Mg^2+^, and K^+^ are depicted in sticks. Water molecules, Mg^2+^ and K^+^ are shown as red, green and purple spheres, respectively. The diameter of the K^+^ ion has been reduced for clarity. Dotted lines denote the octahedral coordination of Mg^2+^ and the distorted polyhedral coordination of K^+^. (**B**) Superimposition of a single mHsp60 subunit from the structures of apo mHsp60_7_ (varying shades of red for equatorial, intermediate and apical domains) and ATP-bound mHsp60_14_ (varying shades of green for each domain). Indicated with arrows and angles are the ATP-triggered rigid body 18.7° counterclockwise of the apical domain and the downward movement of the intermediate domain with the 23.6° downward tilt of helix M. The nucleotide ATP and the Mg^2+^ and K^+^ ions are shown in space-filling representation. (**C**) Side view of two neighboring mHsp60 subunits and close-up of the nucleotide-binding site and intra-ring inter-subunit interface of apo mHsp60_7_ (left, in red) and ATP-bound mHsp60_14_ structures (right, in green). ATP is depicted in sticks, and the Mg^2+^ and K^+^ ions are shown as green and purple spheres, respectively. Helix M, which tilts downwards upon ATP binding, and its interacting residues with neighbor apical or equatorial domain are labeled. Also labeled ß-sheet formed in the intra-ring inter-subunit interface by ß1-ß18 from one subunit and ß2-ß3 from the adjacent subunit.

The binding of ATP triggers conformational changes in each mHsp60 protomer at the level of the intermediate and equatorial domains. Compared to apo mHsp60, in the structure of ATP-bound mHsp60 helix M in the intermediate domain of each subunit tilts down by 23.6° (Fig. 2B; Movie S2). The descent of helix M onto the nucleotide-binding site brings the catalytic residue D399 into contact with the catalytic water and the nucleotide γ-phosphate of ATP (Fig. 2A, 2C right panel; Movie S3). In the equatorial domain (Movie S2) the stem-loop of the ß-hairpin 2/3 flaps upward in the opposite direction to the tilt of helix M (Fig. 2B). As a result, K51 and D52 (both at ß3) contact the nucleotide γ-phosphate and the catalytic water, respectively (Fig. 2A; Movie S3). A 4 Å shift in the loop between helices C/D brings D87 (Fig. 2C), at the top end of helix D, within coordination distance to Mg^2+^ (Fig. 2C; Movie S3). ATP binding also causes the ß-sheet formed by the N-terminal (ß1) and C-terminal (ß18) ß-strands to swing sideways (Fig. 2C; Movie S3).

### Dynamics of the apical domains in ATP-bound mHsp60

In line with the V72I mHsp60 variant^33^, WT mHsp60 also displayed asymmetric apical domain states (Fig. 3). A symmetry expansion, signal subtraction and 3D variability analysis^40^ for ATP-bound mHsp60_14_ identified three classes (Fig. 3A) that show a gradual rigid-body elevation of the apical domain in a sweep arch motion with a maximum tilt of 14.8° (Fig. 3B, left panel; Movie S4).

**Figure 3.**
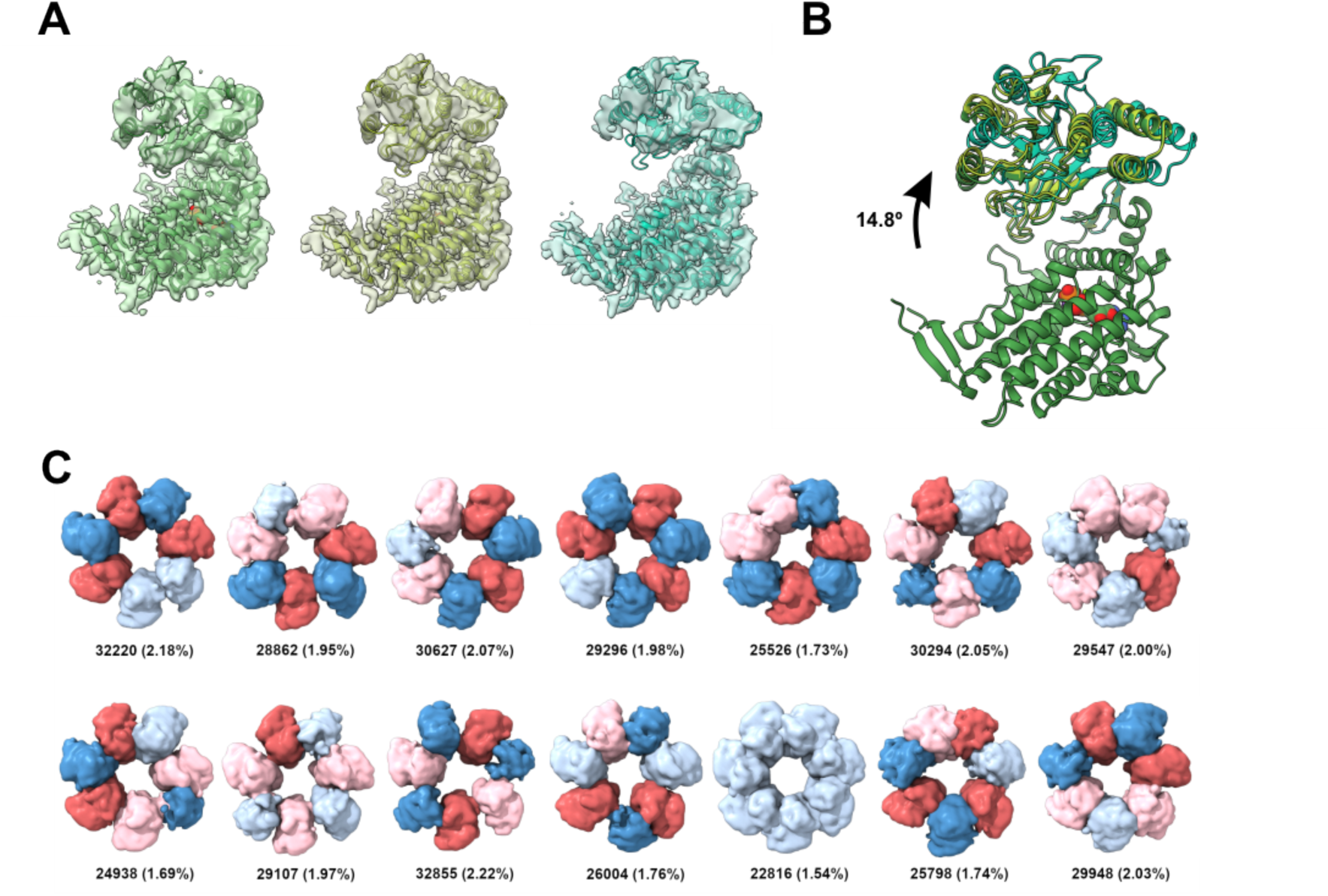
Dynamics of the apical domains upon ATP binding. (**A**) 3D clustered classes resulting from 3D variability analysis (see Methods section) after symmetry expansion and signal subtraction of single subunits of ATP-bound mHsp60_14_. From left to right, the classes show an increasing degree of elevation. For the three classes, a cartoon depiction of the single subunit fitted in the maps obtained is shown. (**B**) Superimposition of the three models derived from the 3D clustered classes as in (A) of ATP-bound mHsp60_14_ obtained by 3D variability analysis depicted in varying shades of green. The classes show a sequential degree of elevation of the apical domains up to 14.8° indicated by an arrow. (**C**) 14 selected classes viewed from the top. Each apical domain is colored based in its position relative to the average: movements towards the protein center are colored in red (0-2 angstroms in pink, and values higher than 2 in deeper red), and movements away are colored in blue (0-2 angstroms in light blue, and more than 2 in dark blue). The number of particles and the percentage for each class are also indicated.

The apical domain rotates 18.7° counterclockwise (view from within the assembly) and rises slightly (Fig. 2B). The position of the apical domain of the second most elevated class corresponds to that observed in the mHsp60_14_ map (Fig. S4, ATP-bound mHsp60_14_ filtered shown in green) used to build the structure (Fig. 1, PDB ID 9ES2). In addition to the subunit-level analysis, we carried out a 3D classification of the apical region at the level of the entire heptameric ring. We symmetry expanded the particles in C7, followed by 3D classification using a soft mask around the apical regions (Fig. 3C). Through this approach, we observed alternate ascending and descending conformations for the apical domain of the mHsp60 subunits similar to those observed for V72I mHsp60^33^. Spatial constraints preclude the possibility of all seven apical domains adopting a downward orientation simultaneously. The up and down movement of the apical domains breaks the 7-fold symmetry of the ring. Thus, WT mHsp60 in the absence of SP displays intrinsic alternating up/down configurations first observed by cryo-EM for the V72I mHsp60 variant in the presence of SP^33^.

### Conformational changes underlying ATP-driven double ring formation

Unlike GroEL, which invariably assembles as a double heptameric ring even without nucleotide^41,42^, mHsp60 can exist as a monomer or assembled as single heptameric rings in the nucleotide-free form, with ATP shifting the equilibrium toward double rings^17,29^. The structures of apo mHsp60_7_ and ATP-bound mHsp60_14_ clarify this behavior.

At the level of the heptameric rings ATP binding induces a major rearrangement of all equatorial domains within each ring (Fig. 4A; Movie S5), including an upward shift of the inter-subunit ß-sheet formed by β-hairpin 2/3 of one equatorial domain and the N- and C-terminal strands (ß1 and ß18) of the preceding subunit (Fig. 4B). This ß-sheet is essential for heptamer assembly^17,18^, and its elevation in the ATP-bound state is linked to interaction of K51 and D52 (strand ß3) with the nucleotide-binding site (Fig. 2A; Fig. 2C, right panel). These rearrangements flatten the equatorial surface of each ring and reorient helix O more parallel to the inter-ring interface. At the same time, helices D and E and their connecting loop are pulled away from the inter-ring interface (Fig. 4A-B). Since the N-terminus of helix D contributes to the nucleotide-binding pocket and its preceding loop engages the nucleotide γ-phosphate, ATP binding directly couples nucleotide recognition to the displacement of helices D and E from the inter-ring interface. A flattened equatorial plane enables the correct inter-ring contacts: a right locus defined by a twofold-symmetrical interaction between opposing S464 residues^17^, and a left locus mediated by weak electrostatic contacts between K109 and E105the formation of the right interacting locus (perspective from inside the assembly) (Fig. 4C). By contrast, in apo mHsp60 the equatorial surface is insufficiently flat, helix O is not parallel to the interface and helices D and E are not pulled away (Fig. S12), explaining why only single rings form in the absence of ATP.

**Figure 4.**
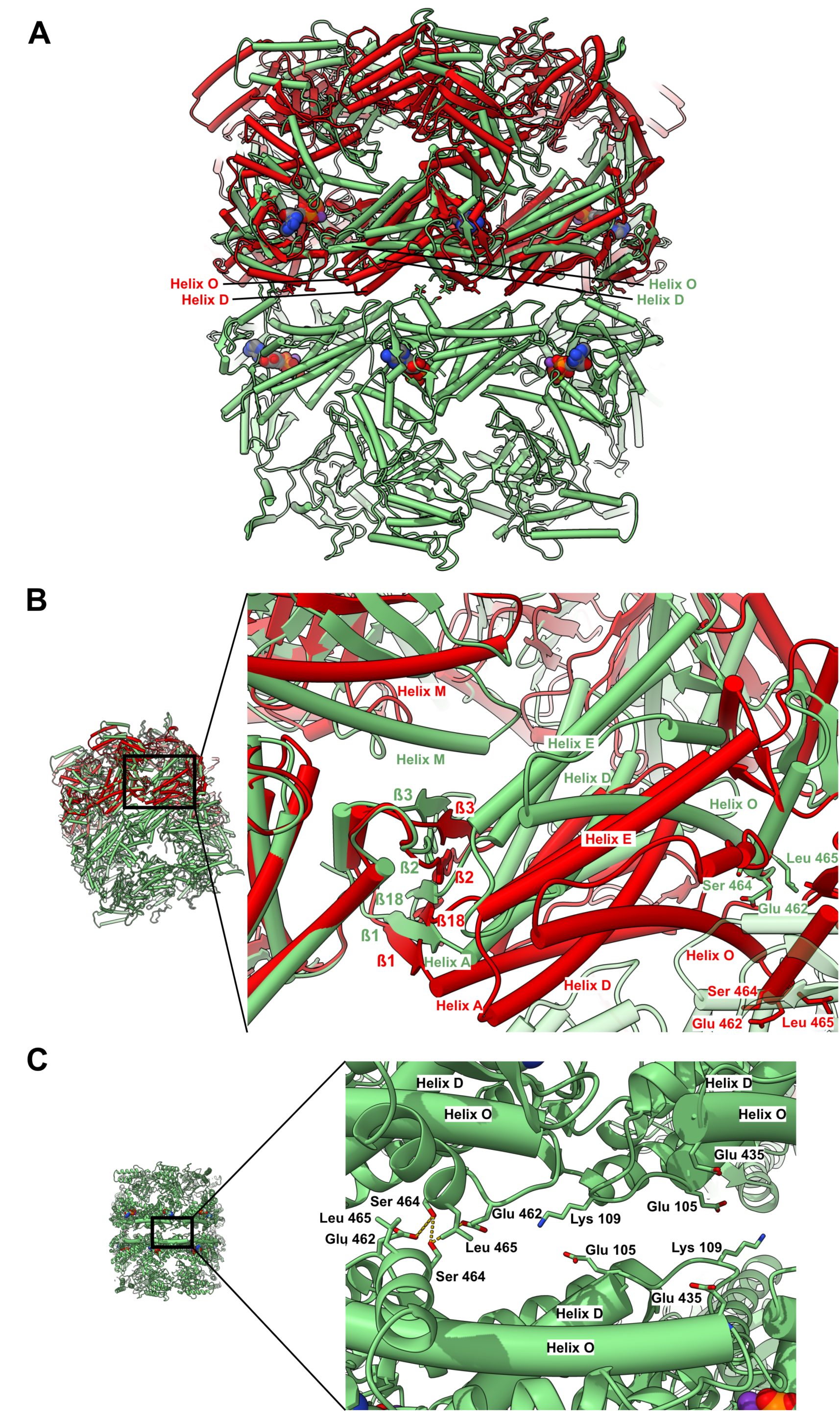
ATP-triggered mHsp60 double-ring formation. (**A**) Side view of a superimposition (without any alignment between subunits) of apo mHsp60_7_ (red) on top of ATP-bound mHsp60_14_ (green). Helices D and O are labelled and the nucleotide ATP and the Mg^2+^ and K^+^ are shown in space-filling representation. Residues involved in inter-ring interactions are shown as sticks for reference. (**B**) Oblique side-view and close-up of a superimposition as in (A), but with a single apo mHsp60_7_ and ATP-bound mHsp60_14_ subunit (on the left) aligned to each other. The ß-strands that form the ß-sheet intra-ring interface and the helices that move when the neighbor subunit is aligned are labelled according to the structure colors. Residues involved in inter-ring interactions are shown in sticks for reference. Subunits across the inter-ring interface in the opposite ring are shown as transparent green. (**C**) Side-view and close-up of selected residues at the double ring interface of ATP-bound mHsp60_14_. Labels mark helices D and O and key residues. Inter-ring interactions are marked with broken lines. ATP is shown in space-filling representation.

### Inner cavity changes between assembly intermediates

Together with the previously reported ADP-bound mHsp60-mHsp10 structures^17^, we now provide a comprehensive set of assembly intermediates elucidated by cryo-EM from the same WT mHsp60 and mHsp10 preparations. At the assembly level, ATP binding to apo mHsp60_7_ induces a radial constriction (12 Å decrease in diameter) of the cavity-facing ring formed by the apical domains (Fig. S13), while the subsequent binding of the heptameric mHsp10 lid causes a radial expansion (32 Å increase in diameter).

Recent cryo-EM evidence suggests the C-terminal 22-amino acid-long tails ending with two GGM repeats, proposed initially to project from the equatorial domains into each ring cavity^17^, are involved in contacts with SP^33^. Although complete GGM-tail density was not visible, we observed an extended density projecting toward the cavity center, with a 5.6 Å reduction in diameter from apo mHsp60₇ to ATP-bound mHsp60₁₄. In apo mHsp60₇, the model extended to residue P526, whereas in ATP-bound mHsp60₁₄ one additional residue (K527) preceding the GGM tail could be modeled. Binding of mHsp10 to ATP-bound mHsp60 also affects the diameter of the cavity-facing ring formed by the equatorial domains. The diameter of this ring does not change significantly in the ATP- and ADP-bound football complexes.

mHsp10 binding completes the formation of a SP folding cage whose inner surface is predominantly hydrophilic and remains largely unaltered upon ATP hydrolysis (Fig. S14). The hydrophobicity of the cavity-facing walls of mHsp60 and GroEL complexes is comparable (Fig. S14), but in contrast to GroEL, the mHsp60 surface is predominantly positively charged (Fig. S15A). Comparison of the ATP-bound mHsp60–mHsp10 football with the ADP:BeF₃-bound GroEL–GroES football (PDB 5OPX) reveals several substitutions of negatively charged GroEL residues by neutral or positively charged residues in mHsp60. These include E63K and E67K in the equatorial domains near the inter-ring interface, E178K and E397T in the intermediate domains, and E232S and E308N in the apical domains close to the mHsp10 interface (Fig. S15B). Equivalent substitutions are conserved across mammalian mitochondrial Hsp60s (Fig. S16). In the co-chaperonin lid, GroES residues I54, E56, and S92 are replaced by K, K, and R in mHsp10, respectively. This distinct electrostatic potential likely contributes to the ability of mitochondrial Hsp60 to accommodate a different spectrum of SPs compared to GroEL^43^.

## DISCUSSION

ATP-driven molecular machines undergo functional cycles through successive transitions between distinct conformational and oligomeric states^44^. Using cryo-EM, we now have a set of high-resolution cryo-EM structures, elucidated from the same recombinant WT mHsp60 and mHsp10 preparations, that delineate how mHsp60 responds to ATP binding and hydrolysis^17^. These structures reveal the conformational changes, domain motions, intra-ring subunit rearrangements and inter-ring contacts that drive double-ring association, thereby defining key intermediates in the mHsp60–mHsp10 reaction cycle (Fig. 5; movies S6-7 summarize the structural and dynamics data presented). This cycle relies on interconnected single- and double-ring folding reaction pathways^17,29,30,32^.

**Figure 5.**
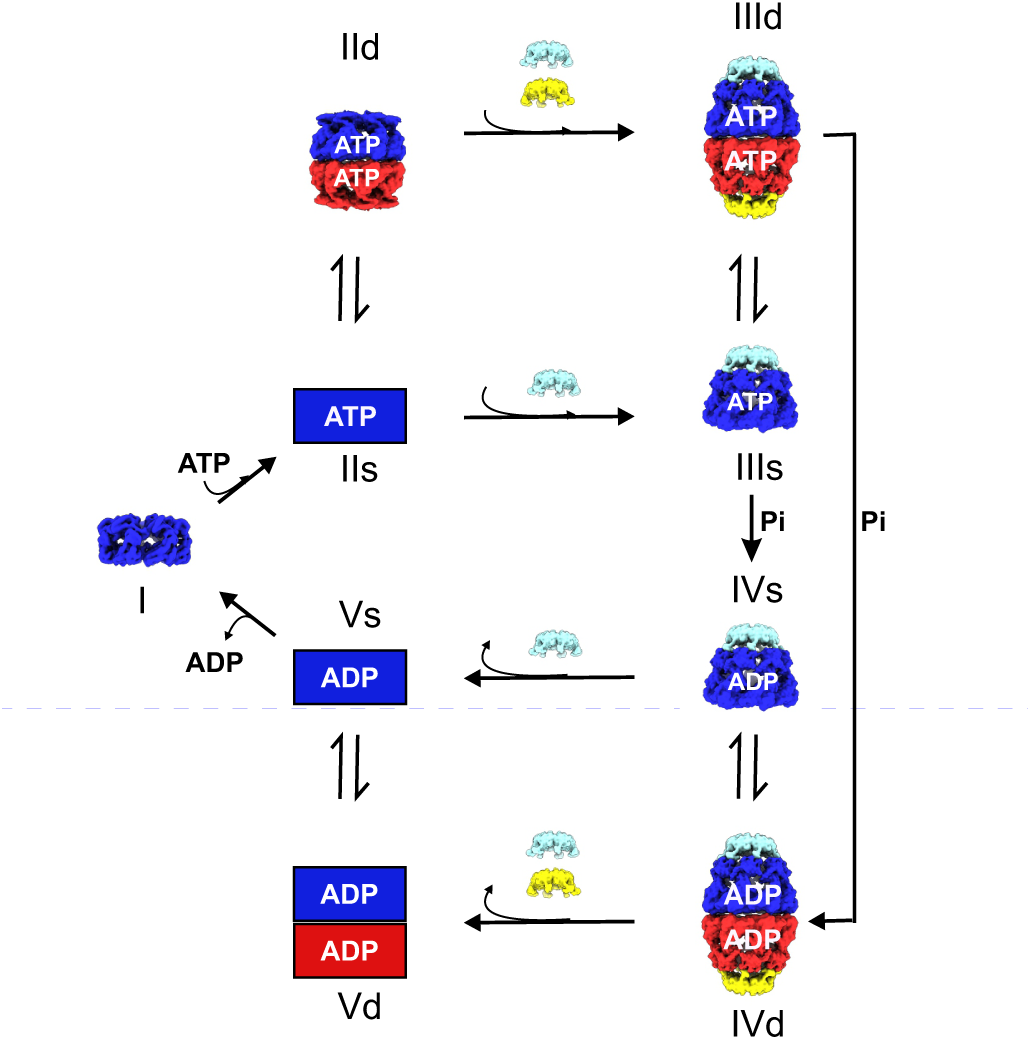
Model for the assembly intermediates of the interconnected single- and double-ring pathways in the reaction cycle of mHsp60:mHsp10. The model depicts ATP binding and hydrolysis driven sequential transitions in the absence of SP based on the *in vitro* cryo-EM structures presented here and from a previous study^17^. Heptameric mHsp60 north and south rings are denoted in blue and red, respectively. Heptameric mHsp10 north and south lids are denoted in cyan and yellow, respectively. Squares represent hypothetical structures. The intermediates are numbered in italics, and s denotes single- and d double-ring complexes. Pi denotes inorganic phosphate.

The question then arises as to how mHsp60 ejects mHsp10. The ATP-triggered descent of the intermediate domain on the nucleotide-binding site, which liberates the apical domain for association with mHsp10, relies on interactions with the γ-phosphate and K^+^ ion with K51, D52, and D399. Hydrolysis of ATP eliminates these interactions (Fig. S11B) and may unlock the intermediate domain and, in a sequence of movements in reverse to those involved in the assembly of mHsp60-mHsp10 complexes, favor the collapse of the apical domains and the ensuing ejection of mHsp10. In such a mechanism, each mHsp60 ring acts as an independent engine, with ATP hydrolysis kinetics triggering each folding chamber to open, in line with reports showing that mHsp60 binds mHsp10 in the presence of ATP, AMPNP and ADP-BeF_3_, but not ADP and ADP-AlF_3_^29,30^. Consistent with mHsp60 nucleotide-dependence studies^29,32^, ADP exchange will occur in the presence of excess ATP (i.e., the 1 mM ATP concentration found in cells). We note that in the reported ADP-bound mHsp60-mHsp10 structures we did not observe mHsp60 rings devoid of co-chaperonin^17^, and we attribute this to the fact that the cryo-EM specimens were vitrified in the presence of low ATP concentration and an excess of mHsp10 over mHsp60, under conditions that do not favor co-chaperonin dissociation.

Thanks to the advent of cryo-EM methods, we unveil here the structural basis for the ATP-triggered assembly of mHsp60-mHsp10 complexes. The structures provide a framework for genetic and biochemical studies to understand the role of mHsp60 in human mitochondrial proteome homeostasis and to discover therapeutics in the treatment of tumors.

## METHODS

### Protein expression and purification

mHsp60 is translated in the cytosol as a 573-amino acid polypeptide containing a 26-amino acid long mitochondrial Matrix Targeting Signal (MTS), which upon cleavage yields a mature 547 amino acid long mitochondrial protein beginning with the sequence AKD (Fig. S1A). The wild-type human mt-cpn60 gene lacking the N-terminal MTS was expressed in *E. coli* with an N-terminal HisTag followed by a TEV cleavage site, and purified by a combination Ni-affinity and anionic-exchange chromatography as previously described^17,38^. As a result of TEV cleavage, the recombinant mHsp60 employed for cryo-EM begins with the amino acid sequence GS. Therefore, the numbering of the recombinant protein begins with G(1)S(2)A(3). Monomeric mHsp60 was concentrated and incubated at 30 °C in the presence of 4 mM ATP, 20 mM KCl, and 20 mM magnesium acetate, in order to induce oligomerization of monomers. After 2 h, the sample was loaded on a Superdex 200 gel-filtration column in 50 mM Tris-HCl pH 7.7, 300 mM NaCl, and 10 mM MgCl_2_. Fractions containing active oligomeric mHsp60 were collected, concentrated, and flash-frozen in liquid nitrogen for storage. Wild-type human mt-cpn10 (Fig. S1B) was expressed without a HisTag in *E. coli* and purified by a combination of anionic- and cationic-exchange chromatography followed by gel-filtration as previously described^17^. Purified mHsp10 in 50 mM Tris-HCl pH 7.7 and 100 mM NaCl was concentrated and flash-frozen in liquid nitrogen for storage.

### Electron microscopy

Pre-incubation of thawed mHsp60 with fresh ATP for 30 min followed by the addition of mHsp10 and then vitrification (within 30 s of the addition of mHsp10) yielded cryo-EM structures of ADP-bound mHsp60-mHsp10 complexes^17^. In the current study, thawed mHsp60 was vitrified without the addition of fresh ATP to obtain the apo form. To prevent ATP hydrolysis and capture ATP-bound mHsp60, thawed mHsp60 was mixed to a final concentration of 20 μM with ATP and immediately vitrified (within ∼10 s of mixing with ATP). To capture ATP-bound mHsp60-mHsp10, thawed 10 μM mHsp60 was mixed in reaction buffer (20 mM Tris-HCl pH 7.7, 20 mM KCl, and 10 mM MgCl_2_) with 2 mM ATP and 12 μM mHsp10 (chaperonin:cochaperonin molar ratio 1:1.2) and immediately vitrified (within ∼10 s of mixing with ATP and mHsp10). For vitrification a 4 μl aliquot of a given mixture was adsorbed onto a glow-discharged UltrAuFoil^®^ R1.2/1.3 (gold foil on gold) 300 mesh grid (Quantifoil), and vitrified in liquid ethane with a Leica GP2 cryoplunger (Leica) using front-side blotting for 2 s at 95% humidity.

Specimens of mHsp60 in the absence or presence of ATP, and of mHsp60-mHsp10 in the presence of ATP were imaged on a 300 kV Krios G4 (ThermoScientific) equipped with a BioContinuum/K3 camera (Gatan) operating in counting mode at a calibrated 0.8238 Å pix^-1^. Employing a 0.8-1.6 μm defocus range we recorded three movies per hole with a total accumulated dose of 50 e^−^ Å^-2^ over 50 frames. Movies were recorded automatically using EPU 2 (ThermoScientific) with Aberration-free image shift (AFIS) and Fringe-free imaging (FFI).

### Cryo-EM data processing and 3D reconstruction

Initial processing of apo mHsp60 and the ATP-bound and mHsp10-bound states followed the same general processing strategy (Fig. S3). Initial frame alignment and contrast transfer function (CTF) estimation was performed using cryoSPARC live^45^. The best movies according to CTF and total motion were chosen for particle selection. Automatic particle picking was performed in cryoSPARC using templates generated from the blob-based picker. Several rounds of reference-free 2D classification were used to clean the data set. Initial models built using *ab initio* reconstruction were used as references for heterogeneous refinement in cryoSPARC to remove poorly aligned particles. Following CTF refinement^46^, 3D homogeneous refinement was carried out with or without imposing symmetry. The local resolution of the 3D reconstructions was determined using cryoSPARC.

To avoid the preferential orientations observed for apo mHsp60_7_, additional 2D classification followed by class rebalance was performed before the last 3D refinement. For ATP-bound mHsp60_14_, after symmetry expansion focused 3D refinement we performed signal subtraction for each particle using a soft mask around a monomer. Subsequently, 3D variability analysis was performed in cryoSPARC^40^. Classes with strong signal for the apical region were selected and refined.

### Apical domain 3D classification

A total of 105.511 aligned particles from the D7-refined ATP-bound mHsp60_14_ were subjected to symmetry expansion. Subsequently, a 3D classification was performed in Cryosparc using a soft mask around the apical regions of one of the rings. For each of the 50 volumes obtained we built a model using the following approach: The consensus model was rigid-body fitted into the volume in ChimeraX^47^. The apical region (residues 190 to 377) of each monomer on the classified layer were also subjected to rigid body fitting. We then calculated the centroid of each apical region and determined its distance to the consensus apical protein center within the x-y plane. Once the models were obtained, we aligned them to attain a low value of the root mean square distance of the apical region to the protein center. We selected the 14 classes in which the apical regions were best resolved.

### Model building

For model building of the apo mHsp60_7_ we began by rigid-body fitting into our map a single mHsp60 subunit (subunit A) of the cryo-EM model of the apo mHsp60 single ring 7AZP^34^. To improve the fit and to optimize stereochemistry, we carried out real space refinement with secondary structure and geometry restraints using Phenix^48^. We used our apo mHsp60_7_ structure (PDB ID 9ES3) as initial model for the ATP-bound mHsp60_14_ map, but each protomer domain (equatorial, intermediate and apical) was rigid-body fitted independently in Chimera^39^. The equatorial and intermediate domains were fitted in the postprocessed map and the apical domain in the Gaussian-filtered map. This model was refined in Phenix^48^ against the postprocessed map, thus the apical region was not refined as the density was very weak. For the football and half-football complexes we used as starting model mHsp60 (subunit A) and mHsp10 (subunit O) of the 3.15 Å crystal structure of the ADP-bound mHsp60^E321K^–mHsp10 football complex (PDB 4PJ1)^18^.

### Figure preparation

All the figures were prepared using CorelDRAW and Adobe Illustrator. All figures containing cryo-EM density maps and protein structures were generated using Chimera^39^.

## SUPPLEMENTARY MATERIALS DESCRIPTION

The supplementary materials for this manuscript includes 16 supplementary figures (Supplementary Figure 1, Supplementary Figure 2, Supplementary Figure 3, Supplementary Figure 4, Supplementary Figure 5, Supplementary Figure 6, Supplementary Figure 7, Supplementary Figure 8, Supplementary Figure 9, Supplementary Figure 10, Supplementary Figure 11, Supplementary Figure 12, Supplementary Figure 13, Supplementary Figure 14, Supplementary Figure 15, and Supplementary Figure 16), 7 supplementary movies (Supplementary Movie 1, Supplementary Movie 2, Supplementary Movie 3, Supplementary Movie 4, Supplementary Movie 5, Supplementary Movie 6, Supplementary Movie 7), and supplementary references.

## ACKNOWLEDGMENTS

We acknowledge support from the United States-Israel Binational Science Foundation (grant number 2015214) to I.U.-B and A.A. I.U.-B was also supported by a grant PID2022-143177NB-I00 from the Spanish State Research Agency and by the Basque Excellence Research Centre program. Y.S. was supported by the European Research Council (ERC) under the European Union’s Horizon 2020 research and innovation programme (grant agreement 723991 - CRYOMATH) and by the NIH/NIGMS Award R01GM136780-01. Negative-stain electron microscopy was carried out at SGIker (UPV/EHU) and initial cryo-EM screening was performed at CICbioGUNE. High-resolution cryo-EM data collection was performed primarily at the Basque Resource for Electron Microscopy located at Instituto Biofisika (UPV/EHU, CSIC), supported primarily by the Department of Education and the Innovation Fund of the Basque Government, with additional support from Fundación Biofísica Bizkaia and MCIN with funding from European Union NextGenerationEU (PRTR-C17.I1). High-resolution cryo-EM data collection was also performed at the Simons Electron Microscopy Center located at the New York Structural Biology Center, supported by grants from the Simons Foundation (SF349247), NYSTAR, and the NIH National Institute of General Medical Sciences (GM103310) with additional support from Agouron Institute (F00316) and NIH (OD019994).

## DATA AVAILABILITY

The EM maps of apo mHsp60, ATP-bound mHsp60, ATP-bound mHsp60-mHsp10 football and half-football complexes have been deposited in the Electron Microscopy Data Bank (http://www.ebi.ac.uk/pdbe/emdb/) under accession numbers EMD-19933, EMD-19932, EMD-19930, and EMD-19931, respectively. The atomic coordinates of apo mHsp60, ATP-bound mHsp60, ATP-bound mHsp60-mHsp10 football and half-football complexes have been deposited in the Protein Data Bank (www.pdb.org) under PDB ID codes 9ES3, 9ES2, 9ES0, 9ES1, respectively. All reagents and relevant data are available from the authors upon request.

## AUTHOR CONTRIBUTIONS

I.T., D.G.-C., J.V., Y. G.-LL., R.M and I.U.-B. performed cryo-EM specimen preparation and data acquisition. I.T., J.P. L.-A., Y. G.-LL., R.M. and Y.S. performed cryo-EM data processing and model building. F.J., M.P., and A.A. performed molecular biology and protein purification. I.T., J.V., A.A., J.A.H. and I.U.-B. wrote the manuscript. I.T., A.A., J.A.H. and I.U.-B. designed the research.

