## Supplementary Information for "Structural basis for ATP-driven double-ring assembly of the human mitochondrial Hsp60 chaperonin"

Igor Tascón, et al.

**This Supplementary Information includes:**

Supplementary Figures 1-16

Legends for Supplementary Movies 1-7

Supplementary References

**Other Supplementary Materials for this manuscript include the following:**

Supplementary Movies 1-7 (mp4 files)

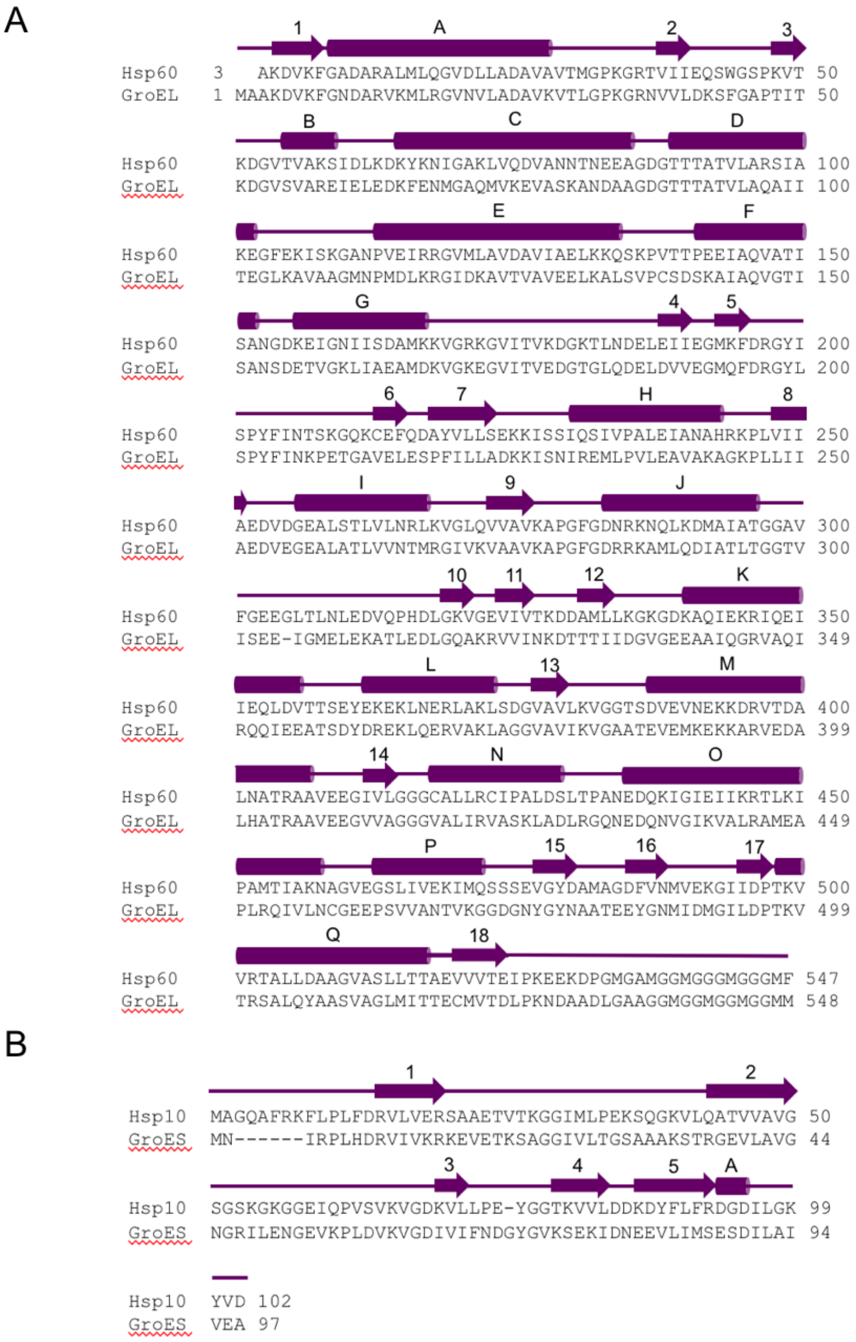


**Supplementary Figure 1. Sequence alignments.** Sequence alignment between (**A**) mHsp60 and GroEL, and (**B**) between mHsp10 and GroES. Secondary structure elements are depicted by purple arrows for β-stands and cylinders for α-helices. The construct for mHsp60 expression contains a TEV cleavage site and upon proteolysis the first two amino acids in mHsp60 are Gly-Ser, which we omitted. The native sequence of mHsp60 begins with Ala at position three.

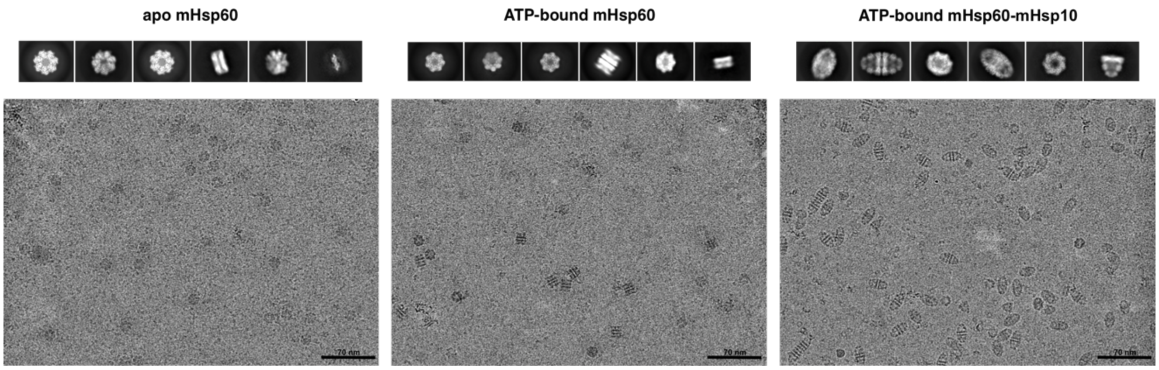


**Supplementary Figure 2. Selected field views.** Selected reference-free 2D class averages and a representative micrograph of the dataset for apo mHsp60, ATP-bound mHsp60 and ATP-bound mHsp60-mHsp10. For apo mHsp60, the classes show front and side views of single-rings, as well as mHsp60 protomers assembled as what appear to be dimers (last class); for ATP-bound mHsp60 the classes show single- and double-rings; and for ATP-bound mHsp60-mHsp10 the classes show football and half-football complexes. In general, single rings can be distinguished from double rings because the latter have D7 symmetry, while the former only display C7 symmetry.

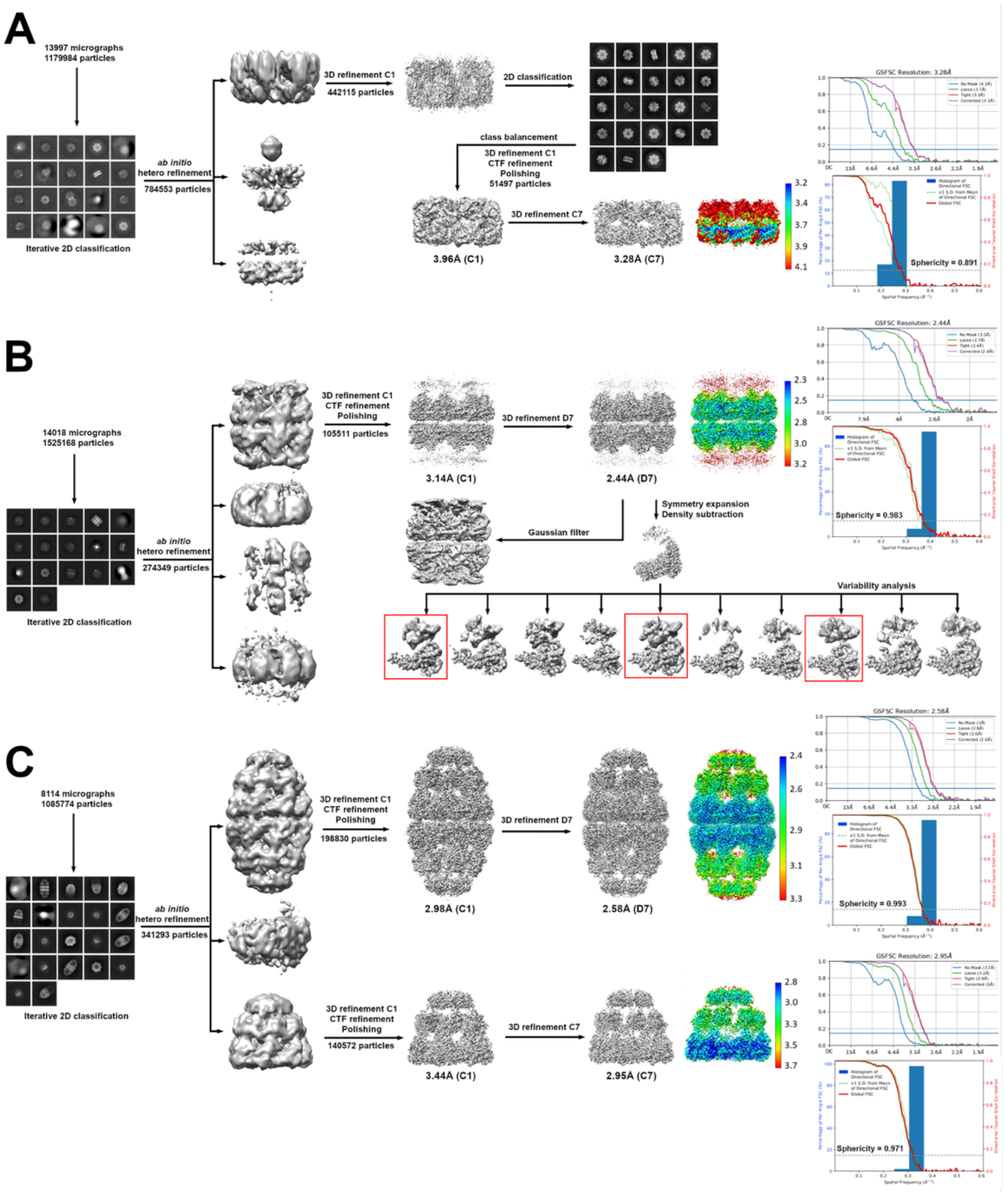


**Supplementary Figure 3. Image processing and 3D reconstruction scheme.** Cryo-EM data processing scheme for (**A**) apo mHsp60$_7$, (**B**) ATP-bound mHsp60$_{14}$ and (**C**) ATP-bound mHsp60$_{14}$-(mHsp10$_7$)$_2$ and mHsp60$_7$-mHsp10$_7$ complexes. Data were processed with cryoSPARC. Processing of all image datasets began with particle picking, iterative 2D classification and ab initio reconstruction. Local resolution and 3D FSC analysis are shown for high-symmetry refinements. After an initial 3D refinement (A) the volume for apo mHsp60$_7$ exhibited a preferential distribution, which was avoided with extra 2D classifications and balancing the resulting 2D classes. In (b) the D7-symmetry refined reconstruction was gaussian-filtered in Chimera using a SD of 1.47 to improve the density of the apical regions. In addition, the data was subjected to symmetry expansion and the signal for all monomers except one was subtracted to each particle. After local refinement, the monomer was subjected to 3D variability analysis in cryoSPARC. The clusters highlighted with a red square contain complete density for the apical region and were used for rigid body fitting (Fig. 3A).

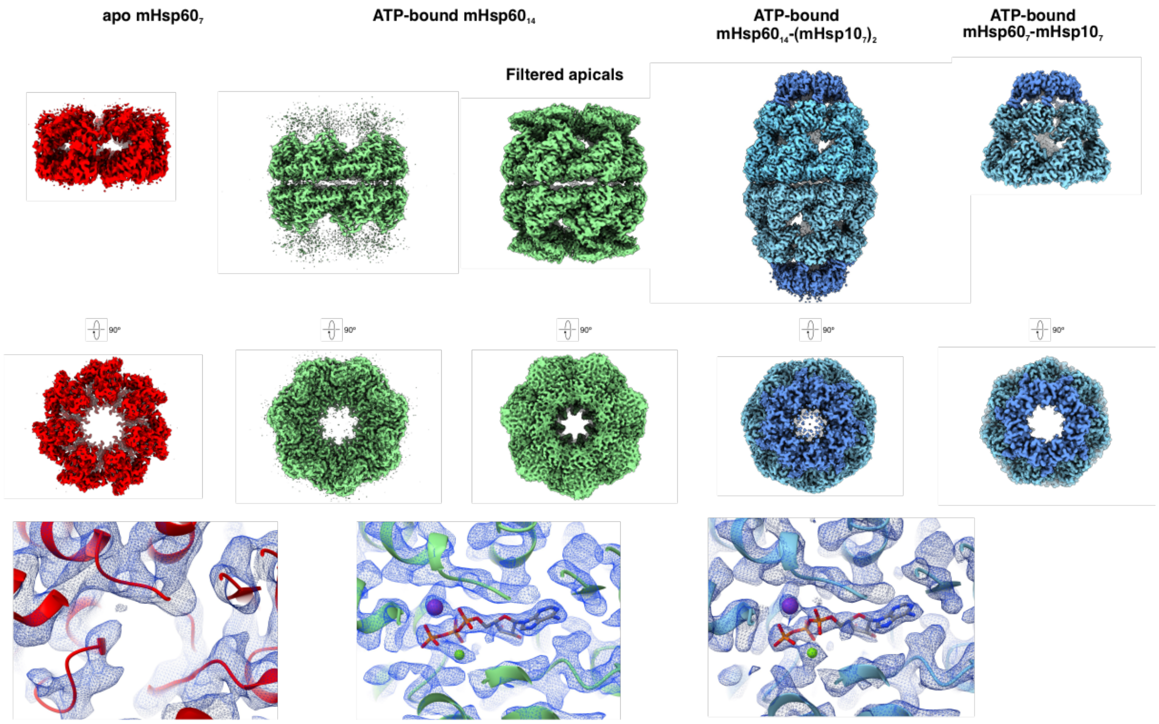


**Supplementary Figure 4. Cryo-EM maps of mHsp60 complexes.** Cryo-EM maps of C7-symmetrized single-ring apo mHsp60$_7$ at 3.28 Å resolution (red), D7-symmetrized double-ring ATP-bound mHsp60$_{14}$ at 2.44 Å resolution (non-filtered and filtered in green), D7-symmetrized double-ring ATP-bound mHsp60$_{14}$-(mHsp10$_7$)$_2$ at 2.58 Å resolution (mHsp60 subunits in cyan and mHsp10 subunits in blue), and C7-symmetrized single-ring ATP-bound mHsp60$_7$-mHsp10$_7$ at 2.95 Å resolution (mHsp60 subunits in cyan and mHsp10 subunits in blue). The maps are shown down a 2-fold dyad orthogonal to the 7-fold molecular axis (side view). Below is a top view down the 7-fold molecular axis. Details of cryo-EM density and atomic model around a single nucleotide-binding site of apo mHsp60$_7$ (red), ATP-bound mHsp60$_{14}$ (green) and ATP-bound mHsp60$_{14}$-(mHsp10$_7$)$_2$ (cyan). ATP is depicted in sticks, and the $Mg^{2+}$ and $K^+$ ions are shown as green and purple spheres, respectively.

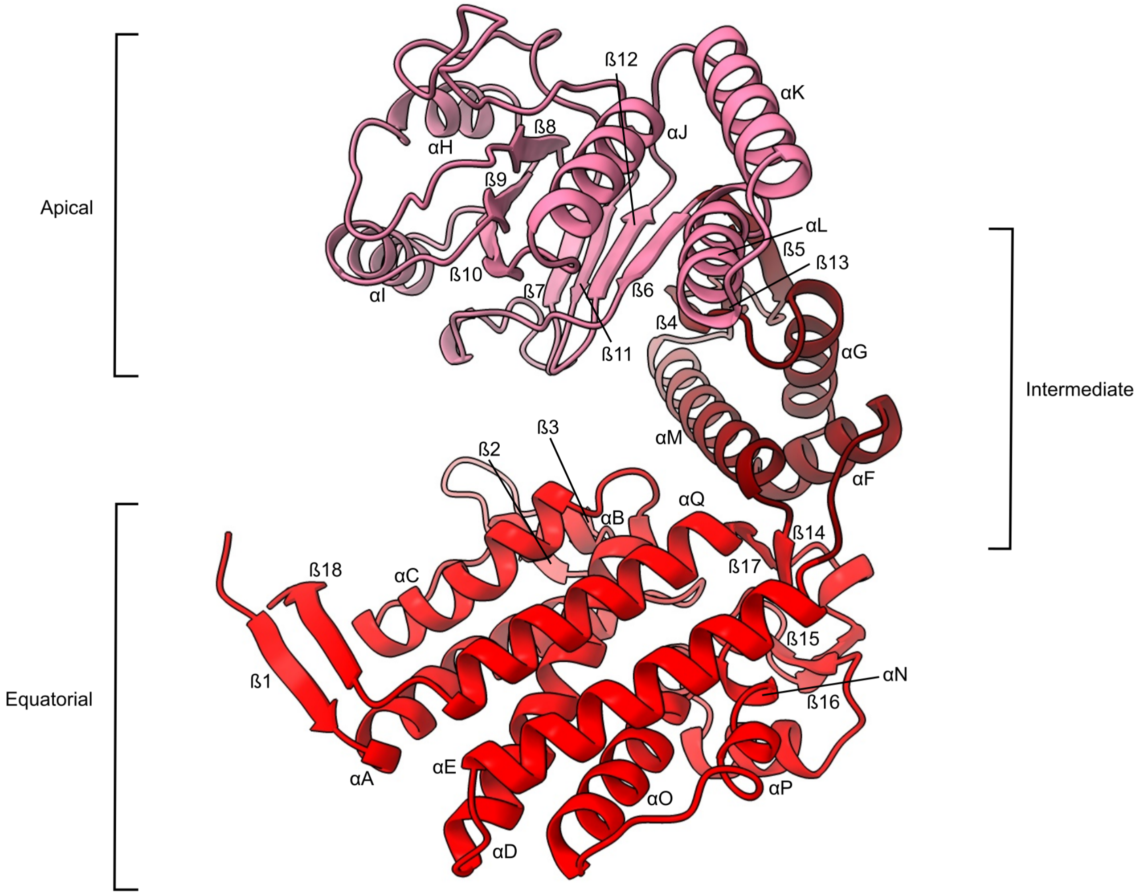


**Supplementary Figure 5. Architecture of mHsp60 subunit.** Side view of one subunit of the structure of apo mHsp60$_7$ showing in red the equatorial domain, in rose the intermediate domain and in pink the apical domain, as well as labelled α-helices and ß-strands.

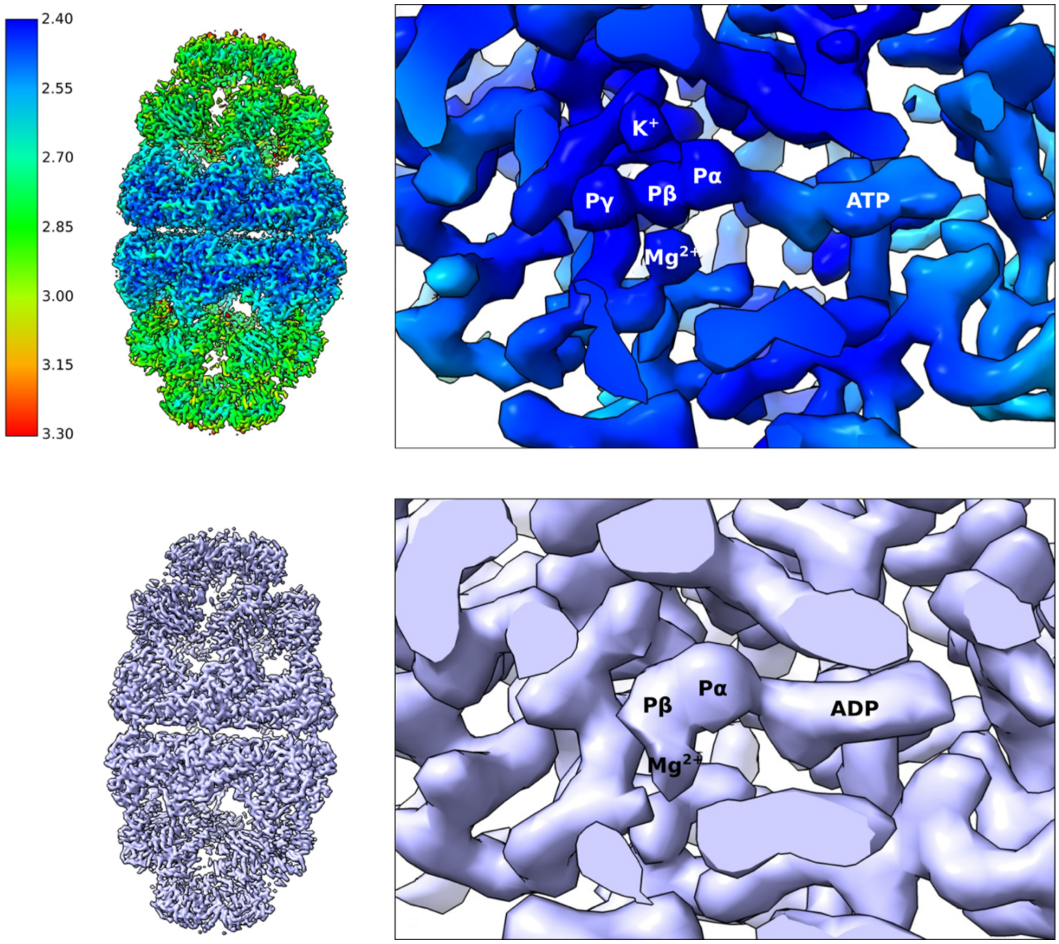


**Supplementary Figure 6. Density for ATP compared to ADP in the nucleotide binding site of mHsp60.** This figure highlights our ability to distinguish ATP from ADP in the cryo-EM maps. The top shows the local resolution by ResMap of the D7 symmetrized 3D reconstruction of ATP-bound mHsp60$_{14}$-(mHsp10$_7$)$_2$ football complex (PDB ID 8B83) determined at 2.58 Å nominal resolution. To the right a zoom in to the densities around the nucleotide binding site, colored in blue by ResMap and at a local resolution of 2.4 Å, of one of the 14 mHsp60 subunits where density details for the adenine base, ribose sugar, and the three serially bonded (α, β, γ) phosphate groups of ATP can be clearly distinguished. In addition, the figure displays clear densities for the $Mg^{2+}$ and $K^+$ ions needed ions for ATP hydrolysis. The bottom shows the cryoEM map of the D7 symmetrized 3D reconstruction of ADP-bound mHsp60$_{14}$-(mHsp10$_7$)$_2$ football complex (PDB ID 6MRC) determined at 3.1 Å nominal resolution (Gomez-Llorente et al 2020). To the right a zoom in to the densities around the nucleotide binding site of one of the 14 mHsp60 subunits where, even at the lower 3.1 Å nominal resolution, density details for just the two (α, β) phosphates of ADP are evident. As expected, here we only see density for $Mg^{2+}$ and not $K^+$.

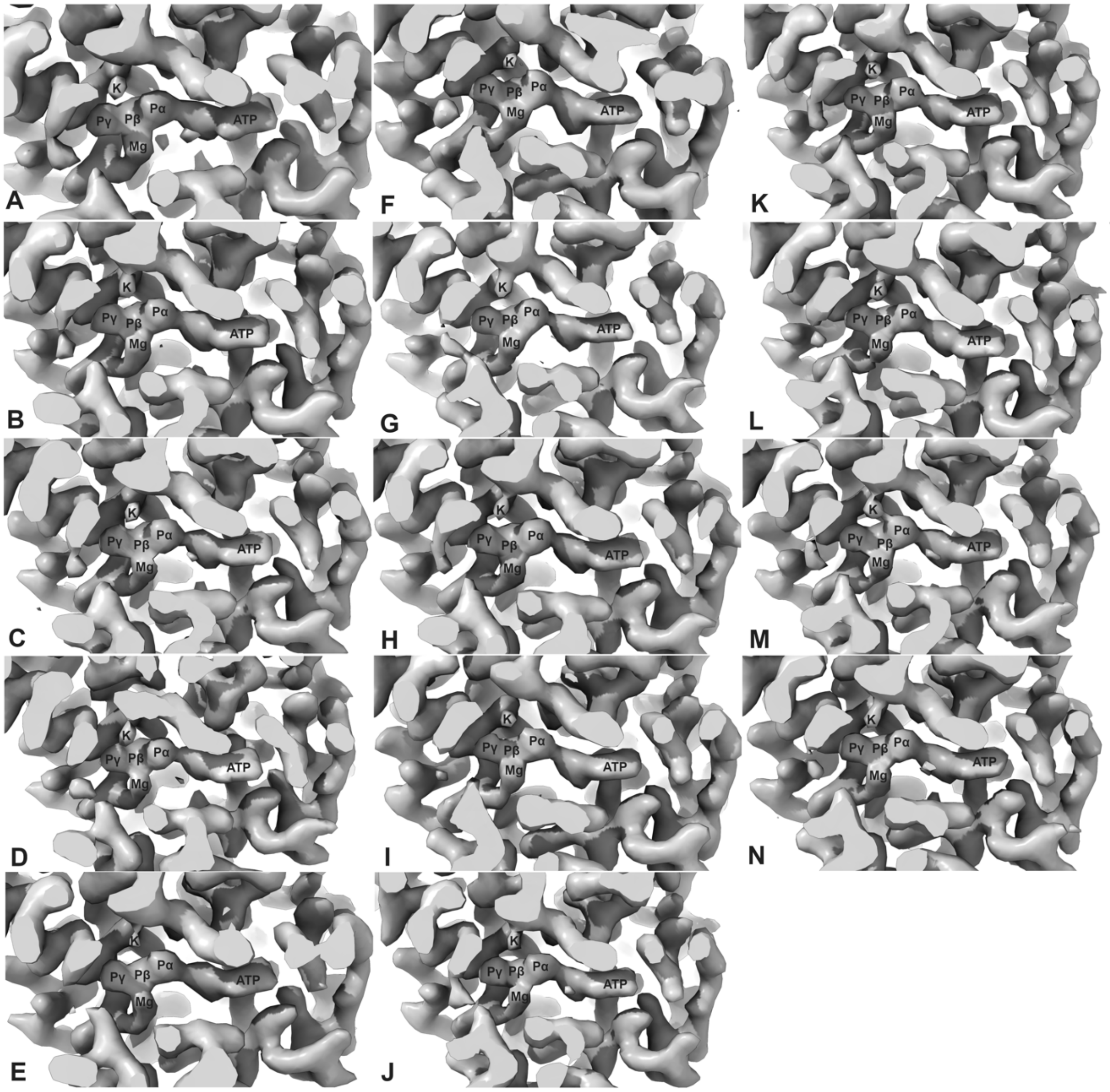


**Supplementary Figure 7. Density for ATP in the nucleotide binding sites of ATP-bound mHsp60$_{14}$-(mHsp10$_7$)$_2$ unsymmetrized (C1) 3D reconstruction.** This figure highlights the occupancy of ATP in all fourteen nucleotide binding sites of a unsymmetrized (C1) 3D reconstruction of mHsp60$_{14}$-(mHsp10$_7$)$_2$ at 2.98Å nominal resolution. The γ-phosphate of ATP can be clearly distinguished in all cases. The densities for the $Mg^{2+}$ and $K^+$ ions needed for ATP hydrolysis are also evident.

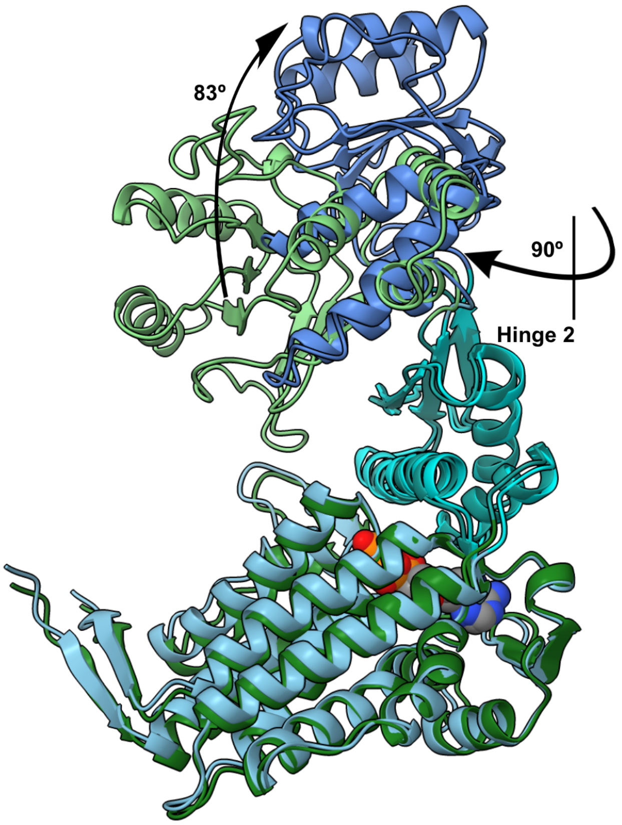


**Supplementary Figure 8. Conformational changes in mHsp60 upon mHsp10 binding.** A superimposition of a single mHsp60 subunit from the structures of ATP-bound $mHsp60_{14}$ (varying shades of green for each domain) and ATP-bound $mHsp60_{14}$-$(mHsp10_7)_2$ (varying shades of blue for each domain). mHsp10 has been removed for clarity in the case of ATP-bound $mHsp60_{14}$-$(mHsp10_7)_2$. Indicated with arrows and angles are the mHsp10-triggered rigid body 90º clockwise rotation and 83º elevation of the apical domain. The nucleotide ATP is shown in space-filling representation.

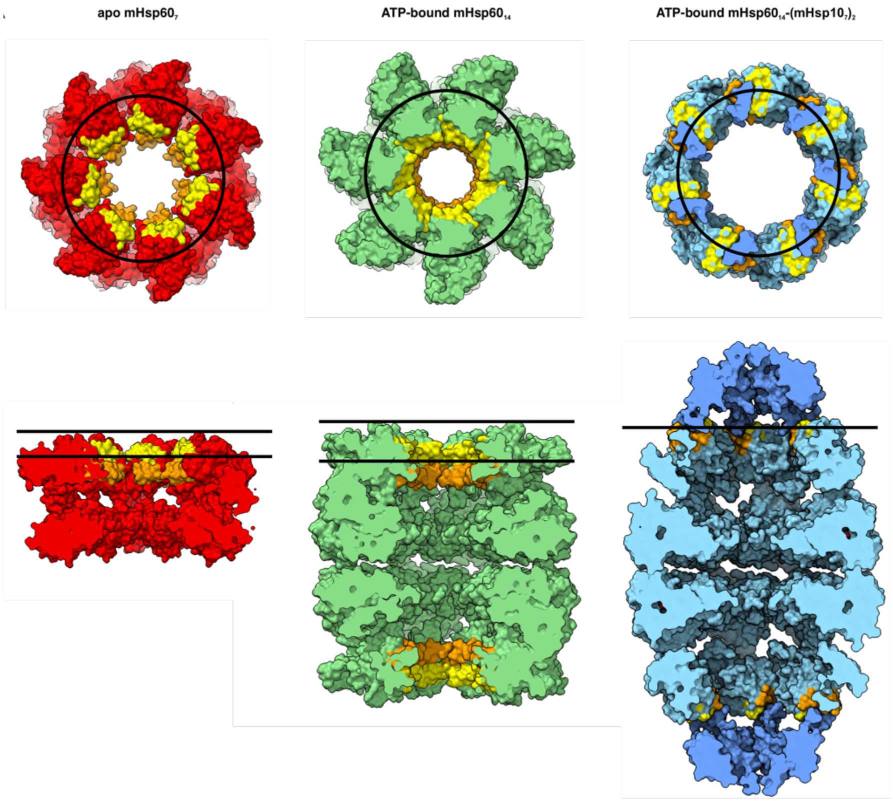


**Supplementary Figure 9. Surfaces in mHsp60 rings that interact with co-chaperonin.** Top views (upper row) and cutaway side views (lower row) of space-filling models of apo mHsp60$_7$ (red), ATP-bound mHsp60$_{14}$ (green), and ATP-bound mHsp60$_{14}$-(mHsp10$_7$)$_2$ (cyan and blue). Polypeptide binding surfaces are colored yellow for helices H and orange for helices I. The interaction sites of mHsp10 mobile loops on the apical domains of mHsp60 are depicted by a black circle. The black lines denote the relative distance between the surface of mHsp60 and the depth of Helix H.

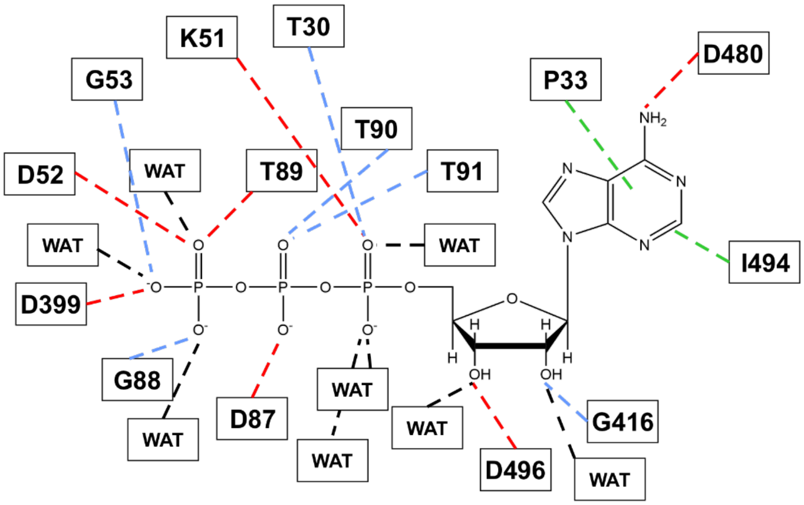


**Supplementary Figure 10. Details of the interactions with ATP.** Interactions between ATP molecule and the surrounding residues and water molecules in the nucleotide binding site of ATP-bound mHsp60$_{14}$. Main chain and side chain interactions are shown with blue and red dotted lines respectively, while π-stacking interactions with green dotted lines. Interactions with water molecules are shown with black dotted lines.

A

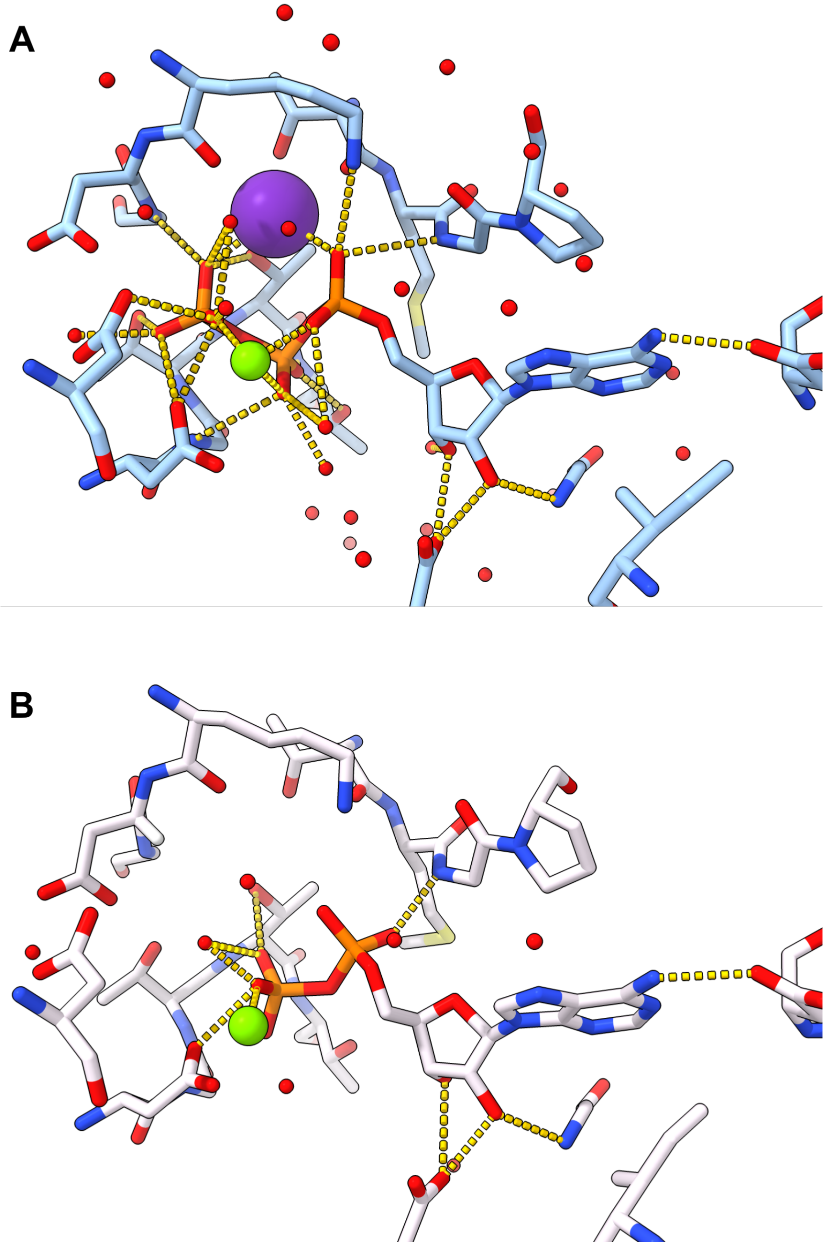

B

**Supplementary Figure 11. Stereochemistry of ATP binding.** (**A**) Stereochemistry of the ATP-bound mHsp60$_{14}$-(mHsp10$_7$)$_2$ nucleotide-binding site. Close-up of the interactions of ATP at the nucleotide binding pocket of a single mHsp60 subunit. Yellow dotted lines denote canonical interactions identified in Coot[54], with hydrogen bonds determined by interaction distance <3.6 Å, and with residues depicted in sticks and the water molecules and $Mg^{2+}$ and $K^+$ ions as red, green and purple spheres, respectively. (**B**) Stereochemistry of the ADP-bound mHsp60$_{14}$-(mHsp10$_7$)$_2$ nucleotide binding site (PDB 6MRC). Close-up of the interactions of ADP at the nucleotide binding pocket of a single mHsp60 subunit. Yellow dotted lines denote the interactions with residues depicted in sticks and the water molecules and the $K^+$ ion as a red and green sphere, respectively.

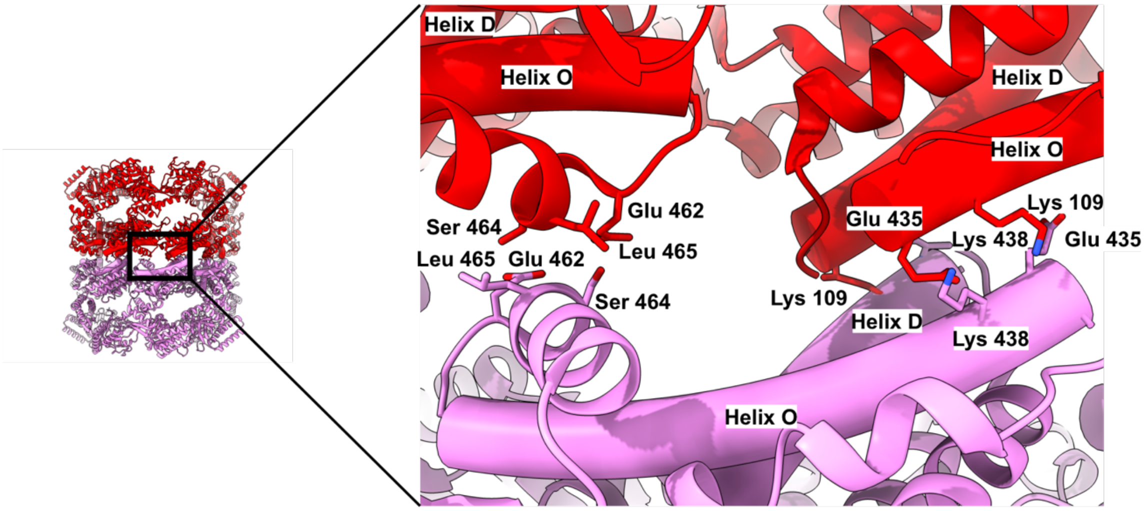


**Supplementary Figure 12. Clashes at the equatorial interface of a hypothetical apo mHsp60$_{14}$ double-ring.** Side-view and close-up of selected residues at the equatorial gap of hypothetical model of apo mHsp60$_{14}$ double-ring (red and pink). The close-up highlights the nonviability of this hypothetical model. A second copy of a single-ring of apo mHsp60$_7$ (pink) has been placed in a position where the interaction of the two rings through E462, S464 and L465 might be plausible. Under such constraints, the clashes between K109 and helix D of opposite rings, or between E435 and K438 across the interface are evident, and would prevent double ring formation. Labels mark helices D and O and key residues.

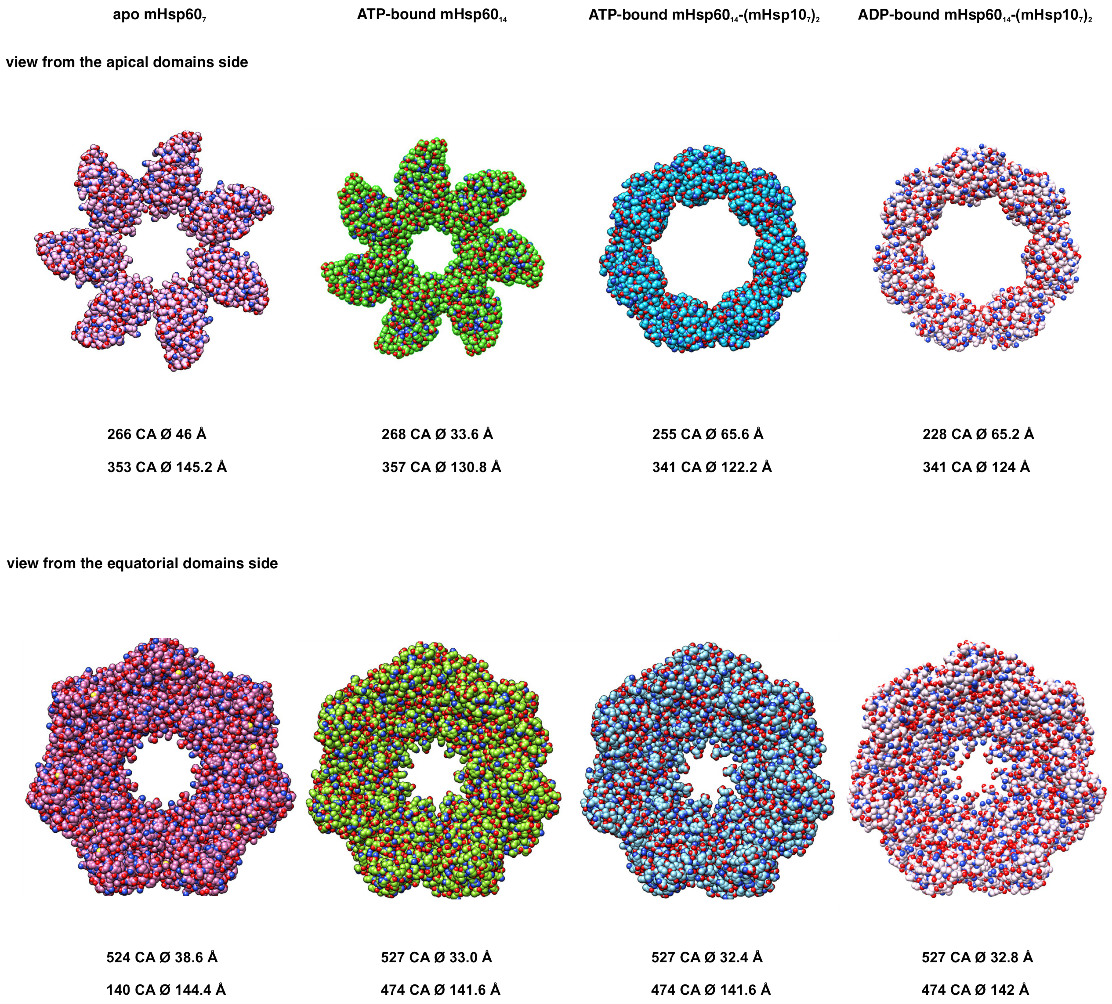


**Supplementary Figure 13. Internal diameter of mHsp60 rings as a function of nucleotide state and mHsp10 binding.** Views of the space-filling representation of apo mHsp60$_7$, ATP-bound mHsp60$_{14}$, ATP-bound mHsp60$_{14}$-(mHsp10$_7$)$_2$, and ADP-bound mHsp60$_{14}$-(mHsp10$_7$)$_2$ (PDB 6MRC). In the top row the views are from the apical domain side (the intermediate and equatorial domains have been removed for clarity), and in the bottom row from the equatorial domain side (the intermediate and apical domains have been removed for clarity). For both views, the shortest diameter and the residues used to calculate it are indicated. Symmetrical Cα were used to define limiting planes.

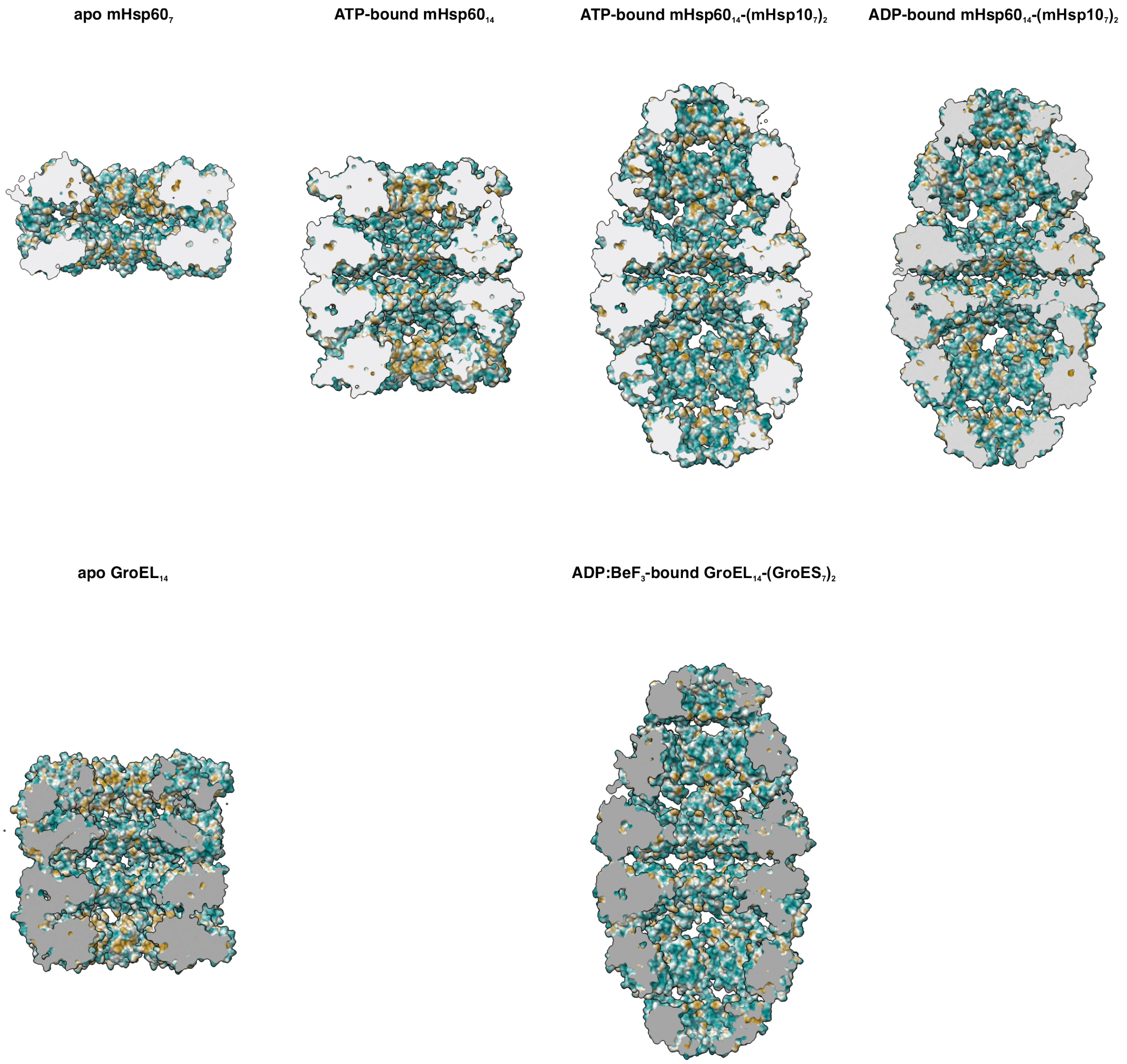


**Supplementary Figure 14. Hydrophobicity of internal surfaces of mHsp60-mHsp10 compared to GroEL-GroES.** The upper row displays central slices parallel to the long axis of apo mHsp60$_7$, ATP-bound mHsp60$_{14}$, ATP-bound mHsp60$_{14}$-(mHsp10$_7$)$_2$, and ADP-bound mHsp60$_{14}$-(mHsp10$_7$)$_2$ (PDB 6MRC) showing the folding cavity in a molecular surface representation colored by hydrophobicity. Dodger blue for the most hydrophilic, to white, to orange for the most hydrophobic. The lower row depicts apo GroEL$_{14}$ (PDB ID 5W0S) and ADP:BeF$_3$-bound GroEL$_{14}$-(GroES$_7$)$_2$ (PDB ID 5OPX).

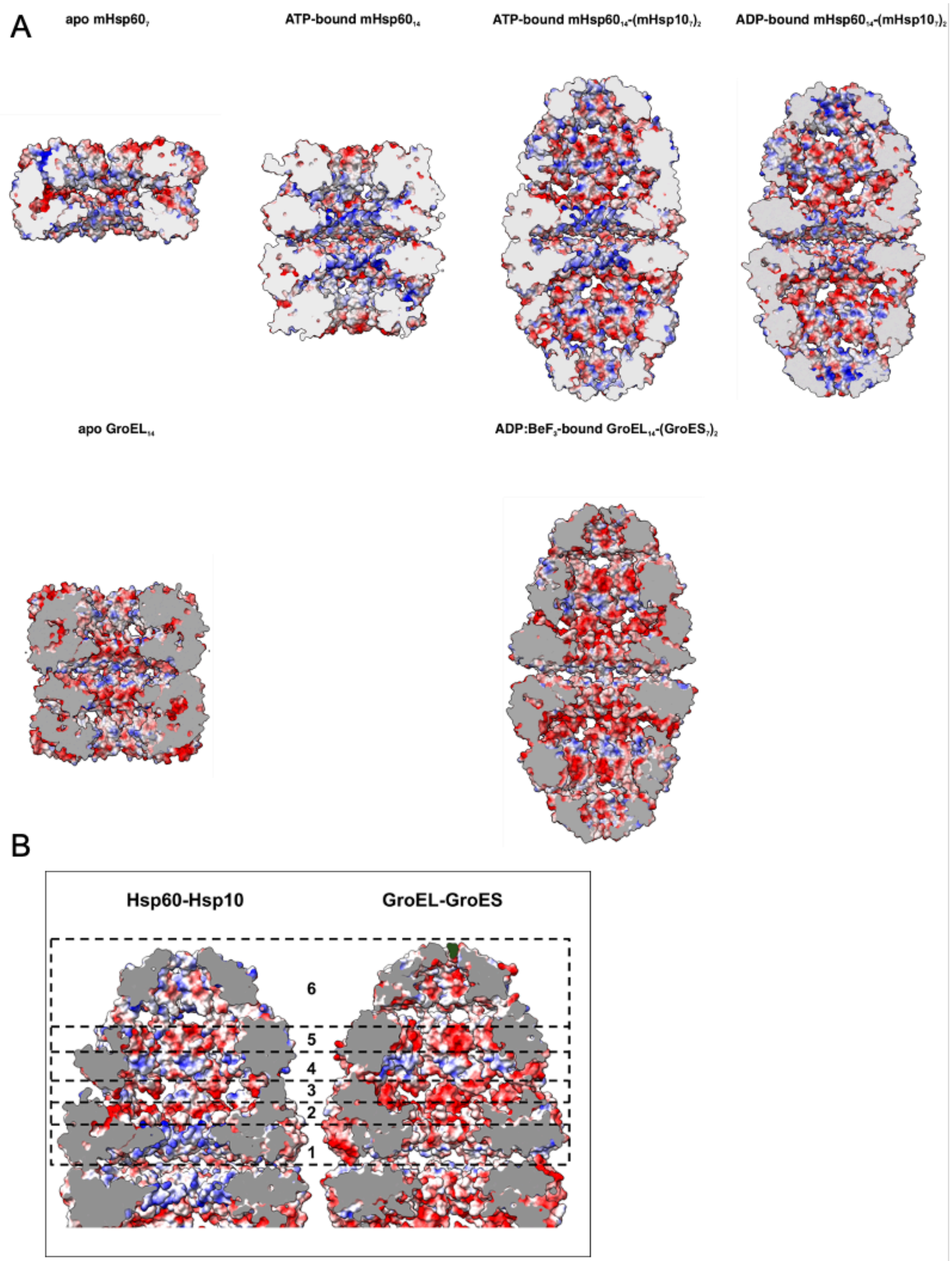


**Supplementary Figure 15. Electrostatics of internal surfaces of mHsp60-mHsp10 compared to GroEL-GroES.** (**A**) The upper row displays central slices parallel to the long axis of apo mHsp60$_7$, ATP-bound mHsp60$_{14}$, ATP-bound mHsp60$_{14}$-(mHsp10$_7$)$_2$, and ADP-bound mHsp60$_{14}$-(mHsp10$_7$)$_2$ (PDB 6MRC) showing the folding cavity in a molecular surface representation colored by electrostatic surface potential. Blue for positive and red for negative charges. The lower row depicts apo GroEL$_{14}$ (PDB ID 5W0S) and ADP:BeF$_3$-bound GroEL$_{14}$-(GroES$_7$)$_2$ (PDB ID 5OPX). (**B**) View of the cavity-facing surface divided in slices perpendicular to the long molecular axis of the football complexes for mHsp60$_{14}$-(mHsp10$_7$)$_2$ (left) and GroEL$_{14}$-(GroES$_7$)$_2$ (right).

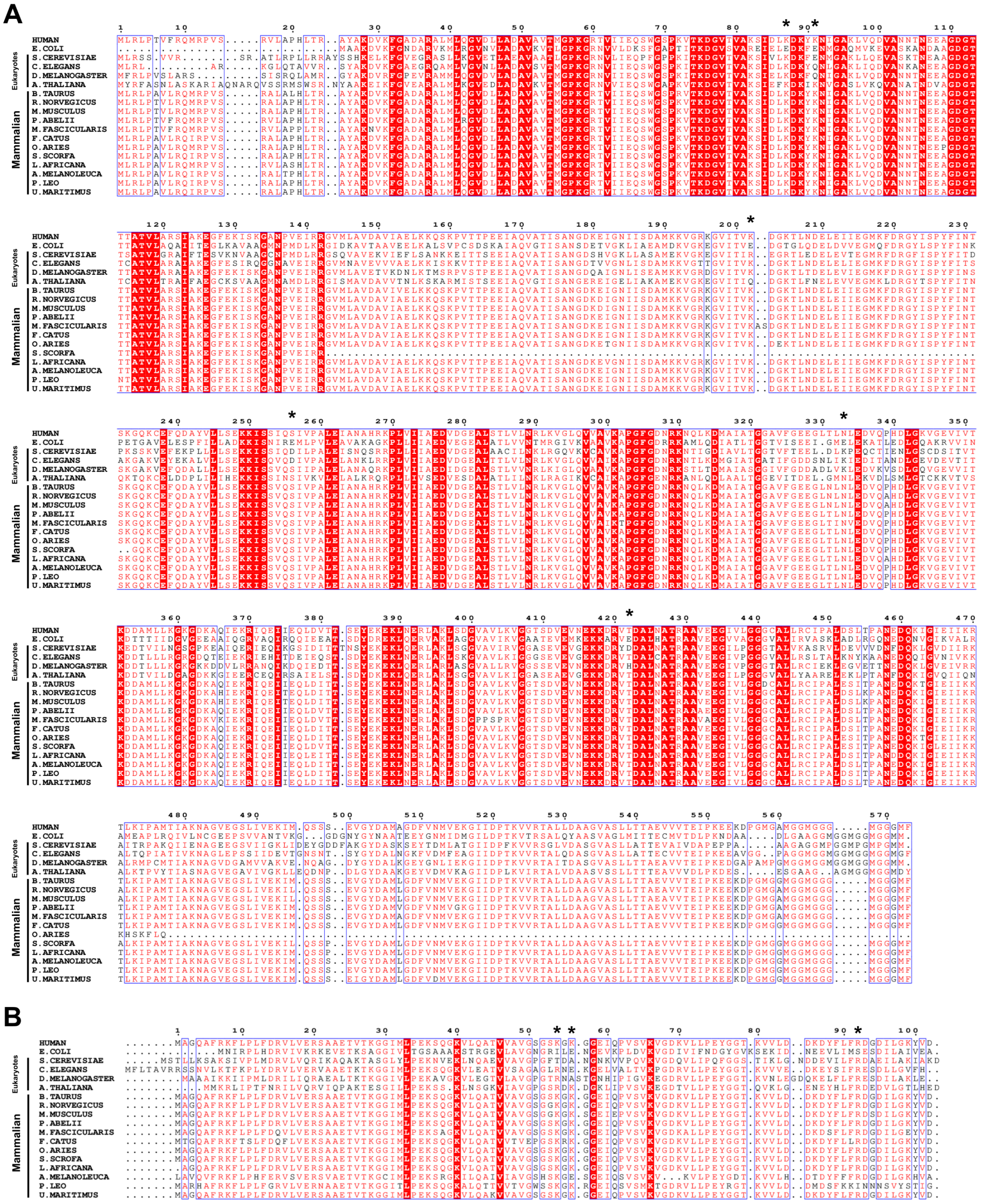

**Supplementary Figure 16. Sequence alignment.** Multiple sequence alignment between human Hsp60 (**A**) and Hsp10 (**B**) and their homologues in mammalian species and selected eukaryotes. GroEL and GroES from E. coli are also included. In the alignment a red box with a white character denotes identity; red character indicates similarity within a group; blue frame indicates similarity across groups; number on top indicates residue number for the first sequence. Selected residues facing the cavity of the chaperonin-cochaperonin complexes are labelled with an asterisk (*****). The multiple sequence alignment was performed in Uniprot (https://www.uniprot.org/align) using 18 entries from UniprotKB corresponding to Hsp60 and Hsp10 homologues. Sequence identity and similarity was analyzed using ESPript3 (https://espript.ibcp.fr/ESPript/cgi-bin/ESPript.cgi).

**Supplementary Movie 1.** Morph between ATP-bound $mHsp60_{14}$ (green) and ATP-bound $mHsp60_{14}$-$(mHsp10_7)_2$ (blue) showing a single subunit. An elevation and rotation of the apical domains is observed. ATP is shown in space-filling representation.

**Supplementary Movie 2.** Morph between a single subunit of apo $mHsp60_7$ (red) and a single subunit of ATP-bound $mHsp60_{14}$ (green). In the intermediate domain, a tilting of down of helix M and the ß-sheet connected to it, and a downward translation of the helices F and G is observed. The movement in the apical domain consists of a counterclockwise rotation with slight elevation. In the equatorial domain, an upwards flapping of the ß-hairpin and the flexing of helix D is shown.

**Supplementary Movie 3.** A rotated close-up view of the morph shown in Movie S2, to highlight the motion of the residues that interact with the ATP. ATP is shown in space filling representation and the residues involved in ATP binding are shown as sticks.

**Supplementary Movie 4.** Morph between the three classes of the single subunits of ATP-bound $mHsp60^{14}$ (varying shades of green) obtained by multi body refinement after symmetry expansion and signal subtraction. A graded elevation throughout the three classes is observed. ATP is shown in space-filling representation.

**Supplementary Movie 5.** Morph between apo $mHsp60_7$ and ATP-bound $mHsp60_{14}$ showing the rearrangement of the heptameric mHsp60 ring to allow double-ring association. ATP is shown in space filling representation.

**Supplementary Movie 6.** Side view of the morph between apo mHsp607, ATP-bound $mHsp60_{14}$ and ATP-bound $mHsp60_{14}$-$(mHsp10_7)_2$ showing full complexes. Rearrangements of the heptameric mHsp60 rings to allow double-ring association and binding of heptameric mHsp10 lids are observed. Heptameric mHsp60 north and south rings are denoted in blue and red, respectively. Heptameric mHsp10 north and south lids are denoted in cyan and yellow, respectively. ATP is shown in space filling representation.

**Supplementary Movie 7.** Top view of the morph shown in Movie S6.
